# Cultural transmission and genomic co-divergence in the willow tit across the Palearctic

**DOI:** 10.64898/2026.04.23.720297

**Authors:** Athena Syarifa, Jochen Martens, Martin Päckert, Laura Kvist, Lei Wu, Yue-Hua Sun, Jochen B. W. Wolf, Ulrich Knief

**Affiliations:** University of Freiburg, Institute of Biology I (Zoology), Evolutionary Biology and Ecology, Hauptstr. 1, DE-79104 Freiburg, Germany; Institute of Organismic and Molecular Evolution (iomE), Johannes Gutenberg University, Mainz, Germany; Senckenberg Natural History Collections, Museum of Zoology, Dresden, Germany; Ecology and Genetics Research Unit, University of Oulu, Oulu, Finland; State Key Laboratory of Animal Biodiversity Conservation and Integrated Pest Management, Institute of Zoology, Chinese Academy of Sciences, Beijing, China; State Key Laboratory of Animal Ecology and Conservation Biology, Institute of Zoology, Chinese Academy of Sciences, Beijing, China; Division of Evolutionary Biology, LMU Munich, Planegg-Martinsried, Germany

**Keywords:** gene-culture coevolution, reproductive isolation, behavioural isolation, learned song, cultural evolution, continental phylogeography

## Abstract

Cultural transmission of mating behaviour can both promote and constrain genetic divergence, yet its long-term population-genetic consequences remain unclear. In songbirds, learned song can generate behavioural isolation, but its potential to shape genome-wide differentiation at continental scales is rarely assessed. Willow tits provide a compelling system, as three culturally transmitted song types are largely allopatric across the Palearctic, while northern populations exhibit mixed repertoires. Here, we combined a chromosome-level reference genome, whole-genome resequencing of 88 willow tits (*Poecile montanus*) spanning all 14 subspecies, and palaeodistribution modelling to reconstruct the species’ evolutionary history across the Palearctic. Phylogenetic and demographic analyses indicate an origin in Asia during the Late Pliocene to Early Pleistocene, followed by expansions that yielded three deeply diverged genomic lineages in the Asian, Central European, and Northern Palearctic regions. The boundaries of these lineages coincide with major song-type divisions. Tests of historical allele sharing show that gene flow occurred preferentially among lineages that share the same or similar song types, even after accounting for geography, consistent with learned song contributing to prezygotic isolation. Peripheral, song-monotypic populations exhibit signatures of repeated bottlenecks associated with glacial isolation, whereas large northern populations retained broader song repertoires and signals of long-term connectivity. These results provide genome-wide continental evidence that culturally transmitted song mirrors and likely reinforces genomic structure through time in a widespread passerine bird.

## Introduction

The relative importance of prezygotic and postzygotic barriers in the speciation process is expected to vary among taxa. In birds, the striking diversity in mating and signalling traits between closely related species indicates that prezygotic barriers may often play a prominent role in speciation (Edwards et al., 2005; Irwin & Price, 1999; Price et al., 2007).

Male bird song encodes species-specific information and functions to attract conspecific females or to defend territories from conspecific males (Catchpole & Slater, 2008). Variation in song can therefore promote assortative mating and, ultimately, reproductive isolation and speciation (Martens, 1996; Slabbekoorn & Smith, 2002; Uy et al., 2018; Whitehead et al., 2019). However, the strength of such isolation, and whether it leads to genetic divergence, depends on how song and mate preferences are transmitted across generations, particularly if these traits are learned (Verzijden et al., 2012).

Among birds, vocal learning is found in three clades: oscine passerines (songbirds), parrots, and hummingbirds (Searcy et al., 2021; ten Cate, 2021). Within songbirds, the extent to which song development depends on learning varies among species, but oscines are generally regarded as vocal learners. Importantly, however, learning systems differ among species: in closed-ended learners, song is acquired during a sensitive phase early in life, whereas open-ended learners can modify their repertoire throughout adulthood (Beecher & Brenowitz, 2005; Brenowitz & Beecher, 2005). Learned song may be transmitted vertically (from parents to offspring), obliquely (from unrelated adults to juveniles), or horizontally (among peers), each affecting the potential for cultural and genetic divergence in distinct ways (Cavalli-Sforza & Feldman, 1981).

When learning is primarily closed-ended and vertical, songs are copied faithfully from parents or nearby relatives, creating a strong association between cultural and genetic inheritance (Yeh, 2019). Such systems promote population-specific song types that reinforce assortative mating and can facilitate genetic divergence (Grant & Grant, 1996, 2010; Kopps et al., 2014; Whitehead, 2017). In contrast, open-ended learning and horizontal or oblique transmission allow individuals to acquire songs from unrelated tutors or across populations (for instance, after dispersal), promoting cultural and genetic homogenization (Kenyon et al., 2017; Secondi et al., 2003).

Beyond sexual selection, song may further be shaped by natural selection: ecological factors such as vegetation structure influence how song transmits through the environment, and selection may favour song characteristics that improve signal transmission between sender and receiver (Boughman, 2002; Derryberry et al., 2018; Dingle et al., 2008; Endler, 1992; Tobias et al., 2010). Morphological adaptations, such as selection on beak shape, can also indirectly affect vocal anatomy and thereby alter song production (Podos, 2001; Podos & Nowicki, 2004; Podos & Schroeder, 2024).

In addition to selection, stochastic cultural processes can shape song evolution. Cultural drift or founder effects may lead to divergence in small or isolated populations (Baker & Jenkins, 1987; Baker, 1996; Parker et al., 2012), and song simplification has been observed when cultural traditions are weak or incomplete (‘withdrawal of learning’) (Thielcke, 1973a). Martens (1996) expanded this concept into an ‘acoustic character shift’ hypothesis, predicting loss of song complexity in sparsely distributed populations with limited cultural transmission—conditions typical for small refugial or island populations. Mechanisms can include founder-induced relaxed sexual selection (Kaneshiro, 1976; Ödeen & Björklund, 2003), less complex songs of dispersing individuals (Bensch et al., 1998), and a lack of tutors during population expansion. Together, these processes illustrate that learned song can either strengthen or erode reproductive barriers, depending on transmission mode, ecological context, and demographic history.

Here, we study the potential role of song in reproductive isolation and lineage divergence in the willow tit (*Poecile montanus*), a small resident songbird distributed across the Palearctic, from Great Britain to eastern Siberia, China, and Japan. Willow tits display three distinct and evolutionarily stable song types (**Fig. 1A**, **Table 1**). The “Alpine” song type consists of a series of mono-frequency whistles, whereas the “Lowland” or “Normal” song type combines frequency-descending elements into syllables (Martens & Nazarenko, 1993). The so-called “Siberian” song is produced by “bilingual” individuals that recognise and sing both Alpine and Lowland song types. Siberian singers thus dispose of a mixed vocal repertoire that comprises two distinct song types. A third, the “Sino-Japanese” song type, also consists of mono-frequency whistles, but these alternate between frequencies (Martens & Nazarenko, 1993). These song types show a largely allopatric distribution (**Fig. 1A**, **Table 1**). Despite the scarcity of behavioural studies in the willow tit, there is evidence from experiments with hand-reared isolated birds that vocal learning plays a major role in song development of tits (Thielcke, 1973b; Yin et al., 2018) and closely related chickadees (Kroodsma et al., 1995; Shackleton & Ratcliffe, 1993).

**Fig 1.**
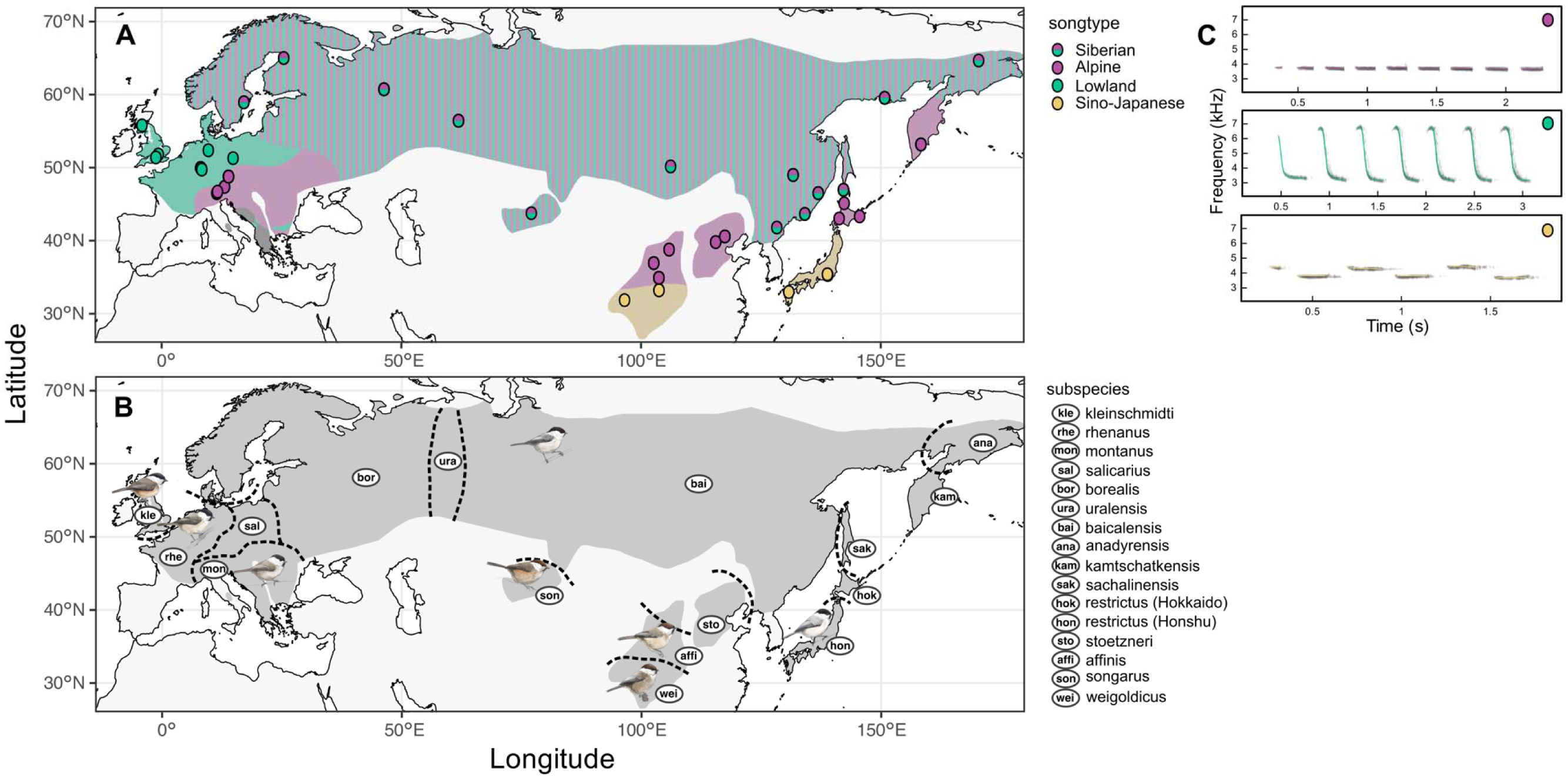
Geographic distribution of cultural and taxonomic variation in the willow tit. (**A**) Distribution of song types across the Palearctic: Lowland (Normal), Alpine, Siberian (mixed Alpine + Lowland), and Sino-Japanese. Dark grey indicates areas with unknown or unassigned song type. The Siberian song repertoire (admixed Alpine and Lowland type) occurs across a broad trans-palearctic range from northern Europe/Scandinavia towards the Russian Far East and Sakhalin. At the eastern and the western margins of the species’ distribution, the Alpine type has a disjunct occurrence, found in the European Alps, Carpathians, and parts of the Balkans, as well as on Hokkaido Island (Japan), in Kamchatka, and in parts of China. The Lowland type occurs in Central Europe and Great Britain, as well as in a disjunct area within the European Balkans. Narrow parapatric contact zones between Alpine and Lowland song types are known from Switzerland and southern Germany, where territorial reactions vary, and boundaries between song types may be either sharp or diffuse (Quaisser & Eck, 2002). At the eastern range margin, the Sino-Japanese type is restricted to parts of China and Honshu Island, Japan (Quaisser & Eck, 2002). Despite their names, no clear ecological correlates explain these distributions: Lowland song also occurs in mountainous areas (e.g. the Rhodopes), whereas Alpine song is also found at low elevations (e.g. in northern Austria). (**B**) Distribution of fourteen recognised subspecies of *Poecile montanus* and *P. weigoldicus*. Polygons represent approximate ranges compiled from Martens & Nazarenko (1993); Martens, Ernst & Petri (1995); Harrap & Quinn (1996); Quaisser & Eck (2002); Thönen (1996). Boundaries, particularly parapatric contact zones, are approximate. (**C**) Representative spectrograms for the Alpine and Lowland song types from personal recordings (U.K.), and for the Sino-Japanese song type from Xeno-Canto (<u>XC145196</u>, Thijs Fijen). Specimen illustrations by Javier Lazaro.

**Table 1.** Subspecies, song type, and geographic occurrence. Siberian song does not represent a distinct song type but a mixed repertoire of both Alpine and Lowland song types.

| Subspecies | Song type | Geographic occurrence |
| --- | --- | --- |
| ***P. montanus*** |  |  |
| *kleinschmidti* | Lowland | Great Britain, restricted to parts of England |
| *rhenanus* | Lowland | Western Europe from North West France, East to Western Germany, and North Switzerland |
| *montanus* | Alpine | Southeast France to Slovakia, Romania, Bulgaria, and North and Central Greece |
| *salicarius* | Lowland | Germany, West Poland, South to North East Switzerland and Austria. Share indefinite boundaries with *montanus* in Northeast Switzerland, Southwest Germany, Northwest Austria |
| *borealis* | Siberian | Fennoscandia, Baltic countries, European Russia, South to West Ukraine |
| *uralensis* | Siberian | Southeast European Russia, West Siberia, and North Kazakhstan. Unclear boundaries with the surrounding subspecies |
| *baicalensis* | Siberian (common), but several songs (Alpine, Lowland, Sino-Japanese) were documented in Altai region (Ernst 1991; Martens, Ernst & Petri 1995) | East Russia (from Yenisei Basin, Altai Mts., East to West coast of Sea of Okhotsk), North Mongolia, Northwest and Northeast China (Xinjiang, northern Inner Mongolia, Heilongjiang, Jilin, and East Liaoning) and North Korea |
| *anadyrensis* | Unknown, but Siberian song was documented in Kolyma Mts. (Quaisser & Eck 2002) | Northeast Siberia, South to North coast of Sea of Okhotsk. Share indefinite boundaries with *baicalensis* in Kolyma Highlands |
| *kamtschatkensis* | Alpine | Kamchatka Peninsula |
| *sachalinensis* | Siberian | Sakhalin Island |
| *restrictus* | Alpine, Sino-Japanese | Hokkaido (Alpine), Honshu (Sino-Japanese), Shikoku, Kyushu |
| *songarus* | Siberian | Southeast Kazakhstan and Kyrgyzstan (Central and East Tien Shan), East to Northwest China (West Xinjiang) |
| *stoetzneri* | Alpine | Northeast China (Southeast Inner Mongolia and Shanxi, East to Beijing, Hebei, and North Henan) |
| *affinis* | Alpine | North Central China (Northeast Qinghai, South Gansu, North Sichuan, Ningxia, and Southwest Shanxi) |
| ***P. weigoldicus*** | Sino-Japanese | South Central China (West Sichuan, Southeast Qinghai, East Xizang, Northwest Yunnan) |

Sources: (Ahmed & Kirwan, 2025; Eck, 1980; Eck & Martens, 2006; Ernst, 1991; Gosler & Clement, 2007; Harrap & Quinn, 1995; Kvist et al., 2001; Martens et al., 1995; Martens & Nazarenko, 1993; Quaisser & Eck, 2002; Thönen, 1962, 1996)

To understand how song variation relates to population history, it is essential to consider the species’ current taxonomic structure. The willow tit currently comprises fourteen recognised subspecies, defined primarily by subtle morphological differences (**Fig. 1B**) (Eck & Martens, 2006; Harrap & Quinn, 1995): *P. m. kleinschmidti*, *rhenanus*, *salicarius*, *montanus* (Great Britain and Central Europe); *borealis*, *uralensis*, *baicalensis* (northern Palearctic, including eastern and northern Europe); *anadyrensis*, *kamtschatkensis* (northeastern Palearctic); *sachalinensis*, *restrictus* (Sakhalin Island and Japan); *songarus* (southern Palearctic); and *affinis*, *stoetzneri* (southeastern Palearctic). Although this morphological classification remains widely used, its delimitation has been inconsistent (Eck, 1980; Eck & Martens, 2006; Gill et al., 2005; Gosler & Clement, 2007; Harrap & Quinn, 1995; Johansson et al., 2013; Quaisser & Eck, 2002; Salzburger et al., 2002; Tritsch et al., 2017). The most taxon-complete molecular phylogeny based on two mitochondrial genes (*ND2*, *COI*) and two nuclear introns (*myo*, *ODC*) supported only three major lineages (Tritsch et al., 2017). A recent genomic study confirmed the genetic distinctiveness of the Chinese subspecies *affinis* and *stoetzneri* relative to their northern Palearctic counterparts (ssp. *baicalensis*) and the narrow-range endemic Sichuan tit, *P. weigoldicus* (Wu et al., 2026).

The recognised subspecies differ slightly in plumage colouration, body size, and relative wing and tail lengths (Eck, 1980; Tritsch et al., 2017). In general, mantle plumage colour varies from yellowish brown in the western part of the range to greyish tones in the east (Eck, 1980; Eck & Martens, 2006). Subspecies in the northern Palearctic tend to have shorter wings and tails compared to those in China, and eastern subspecies and those at higher altitudes are generally larger-bodied than their European and lower altitude counterparts (Bauer, 2013; Eck, 1980). These morphological patterns broadly follow Bergmann’s and Allen’s ecogeographic rules and may therefore be shaped by natural selection in response to ecological factors such as the thermal environment.

While subspecies delineation is based on morphology, song variation provides an independent axis of differentiation, but song type has not been used in subspecies classification. Nevertheless, wherever a song type boundary is present, it typically coincides with a subspecies boundary, though the reverse is not necessarily true (**Fig. 1A, B**, **Table 1**). This suggests that song types are not arbitrarily distributed cultural traits but coincide with morphological variation and reflect deeper lineage splits, possibly shaped by population separation during the last glacial cycles. If learned song has a role in reproductive isolation, we would expect song type and genetic variation to be coupled, implying that song is transmitted across generations in a manner similar to a genetic trait. We would further expect the locations of glacial refugia during the Last Glacial Maximum to align with the present-day distribution of song types.

In this study, we collected 88 willow tit samples across all fourteen subspecies spanning the entire distribution range and encompassing all four song types (Martens et al., 1995; Martens & Nazarenko, 1993; Quaisser & Eck, 2002; Thönen, 1962). We also included three samples of the Sichuan tit (*P. weigoldicus,* hereafter *weigoldicus*), which had previously been recognised as a subspecies and is now separated from the rest of *P. montanus* at the species level (Gill et al., 2025). Finally, we included two marsh tit samples (*P. palustris*), which we used as an outgroup. First, we assembled a high-quality chromosome-level genome of approximately 1,223 Mb in total length, comprising 191 scaffolds, with an N50 of 76 Mb and a largest scaffold of 156 Mb. Then, we conducted whole-genome resequencing of all individuals (∼17.6× coverage) to characterise genetic structure and demographic history. In addition, we compiled species occurrence data to model the ecological niche of the species through time by hindcasting species distribution models onto past climatic and environmental conditions. Using these complementary datasets, we aimed to (i) characterise genome-wide structure and gene flow across the subspecies’ range, (ii) test whether genetic structure aligns with the distribution of song types, (iii) evaluate whether demographic history and glacial refugia predict present-day song diversity and (iv) test whether culturally transmitted song acts as a barrier to gene flow and contributes to lineage divergence.

## Results & Discussion

### Mitochondrial phylogeny and biogeographic origin

First, we reconstructed the phylogeny of all willow tit subspecies and the Sichuan tit from whole mitochondrial genome sequences, using both maximum-likelihood and Bayesian methods (**Fig. 2A**; **Supplementary Fig. S1**). With the Marsh tit as an outgroup, the Sichuan tit (*weigoldicus*) was recovered as the earliest-diverging lineage and used to root subsequent phylogenetic and introgression analyses. The southern and southeastern Palearctic subspecies—*songarus*, *affinis*, and *stoetzneri*—diverged next, consistent with previous studies (Johansson et al., 2013; Tritsch et al., 2017; Wu et al., 2026). Although these lineages are not strictly monophyletic, we hereafter refer to them collectively as the Asian cluster to reflect their shared early divergence and geographic coherence in Asia, where they are geographically separated from the remaining lineages (including *songarus* from the Tien Shan). Within this group, the parapatric southeastern Chinese subspecies *affinis* and *stoetzneri* were not reciprocally monophyletic and are therefore treated as a single group (*affinis* + *stoetzneri*). Together, they formed the earliest diverging lineage (‘China’ in **Fig. 2A**), sister to all remaining willow tit subspecies, while *songarus* diverged next (‘Central Asia’ in **Fig. 2A**).

**Fig. 2.**
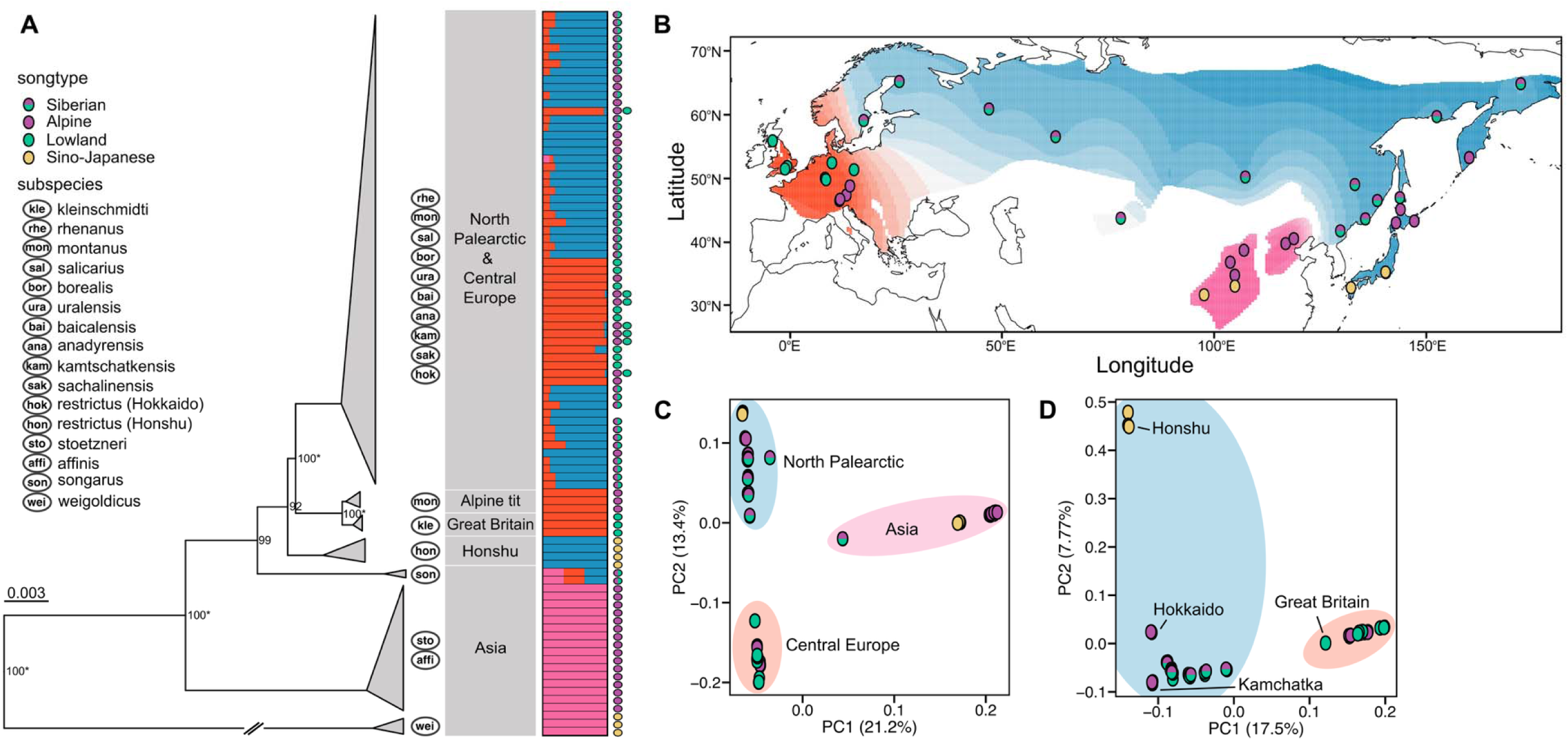
Mitochondrial phylogeny, autosomal ancestry, and genome-wide population structure of the willow tit complex. (**A**) Maximum-likelihood phylogeny based on complete mitochondrial genomes of 91 individuals, rooted with *Poecile palustris* (truncated for space). Branch values represent bootstrap support values > 90 and Bayesian posterior probabilities > 0.99 (*). Scale bar indicates substitutions per site. Subspecies symbols and geographic clusters are denoted on the right of the phylogeny, followed by autosomal ancestry information from ADMIXTURE with K = 3. Symbols denote song types. (**B**) Spatial interpolation of individual ancestry coefficients inferred with *ADMIXTURE* (K = 3) from 1,374,753 autosomal SNPs, showing the three major ancestry groups: Central European (red), Northern Palearctic (blue), and Asian (pink). (**C**) Principal component analysis (PCA) of all willow tit and *P. weigoldicus* individuals (*N* = 91) and (**D**) PCA excluding the three basal Asian lineages (*P. weigoldicus*, (*affinis* + *stoetzneri*), *songarus*; *N* = 72), revealing finer-scale structure within Central European and Northern Palearctic clusters.

These three basal lineages collectively encompass all elements of the three described willow tit song types (Thönen, 1962). The Sichuan tit exhibits the Sino-Japanese song type (Martens et al., 1995; Martens & Nazarenko, 1993; Thönen, 1962), and the southeastern Chinese subspecies *affinis* and *stoetzneri* primarily sing the Alpine song type (Eck & Martens, 2006; Martens et al., 1995). Finally, *songarus* from the Tien Shan mountains displays the Siberian song, which combines Alpine and Lowland song types within individual repertoires (Martens et al., 1995; Martens & Nazarenko, 1993). This pattern is consistent with all song types having been present in the common ancestor of the species, followed by lineage-specific loss, although alternative explanations, such as convergent evolution or cultural exchange, remain alternatives.

Among the remaining taxa, several additional monophyletic groups corresponded to previously described subspecies. The Japanese subspecies *restrictus* diverged next (**Fig. 2A**). Notably, the monophyletic group included only individuals from the southern main island, Honshu, and not from the northern island, Hokkaido. This geographic separation is also reflected in song types, with birds on Honshu singing the Sino-Japanese type and those on Hokkaido the Alpine type. The boundary between these two islands coincides with a well-known biogeographic divide known as Blakiston’s Line, where the Tsugaru Strait (130–140 m deep, 24–40 km wide) serves as a barrier to the movement of terrestrial species (Blakiston, 1883; Dobson, 1994; McKay, 2012). It has also been described as an effective barrier to song in other birds [goldcrest (*Regulus regulus*)] (Martens et al., 1998). During the Last Glacial Maximum (26,000–18,000 years ago) and previous ice ages, Honshu was connected to the Asian mainland via the Korean Peninsula, whereas Hokkaido was connected to northeastern Siberia and Sakhalin. This historical separation most likely underlies the observed divergence in both genetic ancestry and song type between the two *restrictus* populations, especially given that the Sino-Japanese song type is otherwise only found in the southeastern Sichuan tit, whereas the Alpine song type is the exclusive type on Kamchatka. To acknowledge the deep phylogenetic split, we hereafter refer to the two *restrictus* populations on Honshu and Hokkaido as *restrictus* (Honshu) and *restrictus* (Hokkaido), respectively.

The final well-supported split in the mitochondrial phylogeny separated a clade comprising *kleinschmidti* (‘Great Britain’ in **Fig. 2A**) and some *montanus* individuals (‘Alpine tit’ in **Fig. 2A**) from all remaining individuals, including other *montanus* from the same region and individuals from across the northern and northeastern Palearctic, Sakhalin, and Hokkaido (**Fig. 2A**).

The basal position of the southern and southeastern Palearctic subspecies and the localised diversity of song types indicate that the willow tit likely originated in Asia, from where it subsequently expanded across the Palearctic. Previous studies support a Chinese origin of the genus *Poecile*, and of the willow tit in particular (Johansson et al., 2018; Tietze & Borthakur, 2012). Peripheral island populations—on Honshu, Great Britain, and partly in the European Alps—were likely isolated and may have diverged further through genetic drift, whereas the more widespread northern Palearctic subspecies remained connected by gene flow. In particular, populations in Great Britain and the European Alps may have been part of an ancestral population in Central Europe during the Last Glacial Maxima, and subsequently separated during warming events.

By using whole-mitochondrial genomes, we resolved more of the fourteen recognised subspecies than previous molecular studies, which had grouped all but the three most divergent basal lineages in our study—*weigoldicus*, (*affinis* + *stoetzneri*), *songarus*—into a single clade (Tritsch et al., 2017). To further increase phylogenetic resolution and assess genome-wide concordance, we next turned to nuclear whole-genome data.

### Autosomal phylogenetic structure and admixture

To complement the mitochondrial phylogeny, we analysed whole-genome autosomal data after removing multiallelic sites, linked sites (r^2^ > 0.1), and low-complexity regions. The filtered dataset comprised of 1,374,753 SNPs. First, these variants were used to reconstruct maximum likelihood gene trees for 50kb windows sampled every 1Mb along the genome, and the resulting trees were used to construct a coalescence-based species tree with quartet branch support (**Supplementary Fig. S2**). Then, we conducted principal component analyses (PCA) and estimated individual ancestry proportions with the model-based program ADMIXTURE across K = 2–8 genetic clusters (**Supplementary Fig. S2**). Including all fourteen willow tit subspecies and the Sichuan tit (*weigoldicus*), the cross-validation error was lowest at K = 3, which we therefore selected as the best-fit model (**Fig. 2A**, **B**, **Supplementary Fig. S3**). PCA including all individuals yielded PC1 explaining 21.2% and PC2 13.4% of the total variance (**Fig. 2C**). After excluding the Asian cluster (*weigoldicus*, (*affinis* + *stoetzneri*), *songarus*), PC1 and PC2 explained 17.5% and 7.8% of the variation, respectively (**Fig. 2D**).

In the best-fit *ADMIXTURE* model (K = 3), two of the three basal lineages—*weigoldicus*, (*affinis* + *stoetzneri*)—formed a single genetic cluster (**Fig. 2A**, **B**). *Songarus* showed mixed ancestry between this group and the others (**Fig. 2A**), a pattern we interpret cautiously, given its low nucleotide diversity and strong drift signal (see below). It resolved into a distinct cluster at K ≥ 4 (**Supplementary Fig. S2**). Subspecies from Great Britain and Central Europe (*kleinschmidti*, *rhenanus*, *salicarius*, *montanus*) formed a second, separate cluster (**Fig. 2A**, **B**). The remaining subspecies from the northern and northeastern Palearctic—*borealis*, *uralensis*, *baicalensis*, *anadyrensis*, *kamtschatkensis*, *sachalinensis*, and *restrictus* (Honshu and Hokkaido)*—*formed the third cluster (**Fig. 2A**, **B**). The clustering thus broadly reflected the geographic distribution of the subspecies, and we hereafter refer to them as the Asian, Central European, and Northern Palearctic clusters (**Fig. 2B**). The PCA and the autosomal phylogeny supported the *ADMIXTURE* results and confirmed the same three major clusters, with *songarus* positioned centrally between them in the PC1-PC2 plot (**Fig. 2C**).

To evaluate whether this structure reflects continuous isolation-by-distance or localised barriers to gene flow, we estimated spatially explicit migration surfaces using FEEMS (Marcus et al., 2021). FEEMS models spatially heterogeneous isolation-by-distance and identifies regions of reduced effective migration consistent with potential barriers to gene flow. The inferred migration surface supported the three major genomic clusters and revealed pronounced barriers separating the Asian, Central European, and Northern Palearctic clusters, as well as additional barriers at geographic margins (**Supplementary Fig. S4**). In particular, the Sea of Japan emerged as a prominent barrier between Japanese populations and the mainland, consistent with the distinct ancestry of *restrictus* populations on Honshu and Hokkaido.

Although clusters were clearly distinct, particularly in the PCA, a modest degree of admixture persisted between the Central European and Northern Palearctic clusters, fading gradually with increasing distance from their shared boundary. *Salicarius*, located at the eastern edge of the Central European cluster, showed traces of Northern Palearctic ancestry, whereas several Northern Palearctic subspecies (*borealis*, *uralensis*, *baicalensis*, *sachalinensis*, and *anadyrensis*) carried some Central European genetic ancestry (**Fig. 2A**, **Supplementary Fig. S2**, K = 3). These patterns suggest that, following an initial colonization of Central Europe and the Northern Palearctic from an Asian origin, the three major genetic clusters became geographically and genetically isolated—possibly during the Last Glacial Maximum or earlier glacial cycles—and diverged. Secondary contact later occurred between the Central European and Northern Palearctic clusters, resulting in limited gene flow along their shared boundary, while no evidence of gene flow was detected between these two clusters and the allopatric Asian subspecies.

Starting at K = 5, the three basal lineages—*weigoldicus*, (*affinis* + *stoetzneri*), *songarus*—each showed distinct ancestries. At K ≥ 6, *restrictus* (Honshu) also formed a separate ancestry group. These separations were consistent across analyses, matching both the mitochondrial phylogeny and the autosomal PCA (**Fig. 2A, D**).

At K = 8, finer-scale population structure became evident within both the Central European and the Northern Palearctic clusters, consistent with the PCA performed after excluding the Asian cluster (**Supplementary Fig. S2**, K = 8, **Fig. 2D**). PC1 separated the two major clusters, Central European and Northern Palearctic, with subspecies positioned along this axis according to their geographic distances. PC2 distinguished *restrictus* (Honshu) from the remaining Northern Palearctic cluster. This pattern was again consistent with the mitochondrial phylogeny, which likewise indicated distinct ancestry for *restrictus* (Honshu).

Overall, the three major genomic clusters—Asian, Central European, and Northern Palearctic—indicate long-term regional separation. Within these clusters, genetic differentiation followed geography and was consistent with isolation-by-distance, whereas peripheral populations were disproportionately differentiated. The geographic arrangement of song types broadly matched this genetic structure, with song-type boundaries coinciding with major genetic breaks (**Fig. 1A**, **Fig. 2**)

### Historical gene flow and song-associated barriers among willow tit lineages

If historical admixture occurred between the three major ancestry clusters, the phylogenetic position of admixed taxa such as *songarus* would not be well represented in a bifurcating tree. Alternatively, demographic processes such as recent bottlenecks could produce similar signals (Lawson et al., 2018). To disentangle these scenarios, we inferred population splits and migration events using TreeMix (Pickrell & Pritchard, 2012), and validated all inferred migration edges with Patterson’s D-statistics and f_4_-ratio tests. We further visualised excess allele sharing as f-branch statistics (ƒ_b_(C); **Fig. 3A**) (Patterson et al., 2012). All inferred migration edges were supported by D-statistics and f-branch tests (**Supplementary Table S1**), confirming genuine gene flow rather than stochastic drift.

**Fig. 3.**
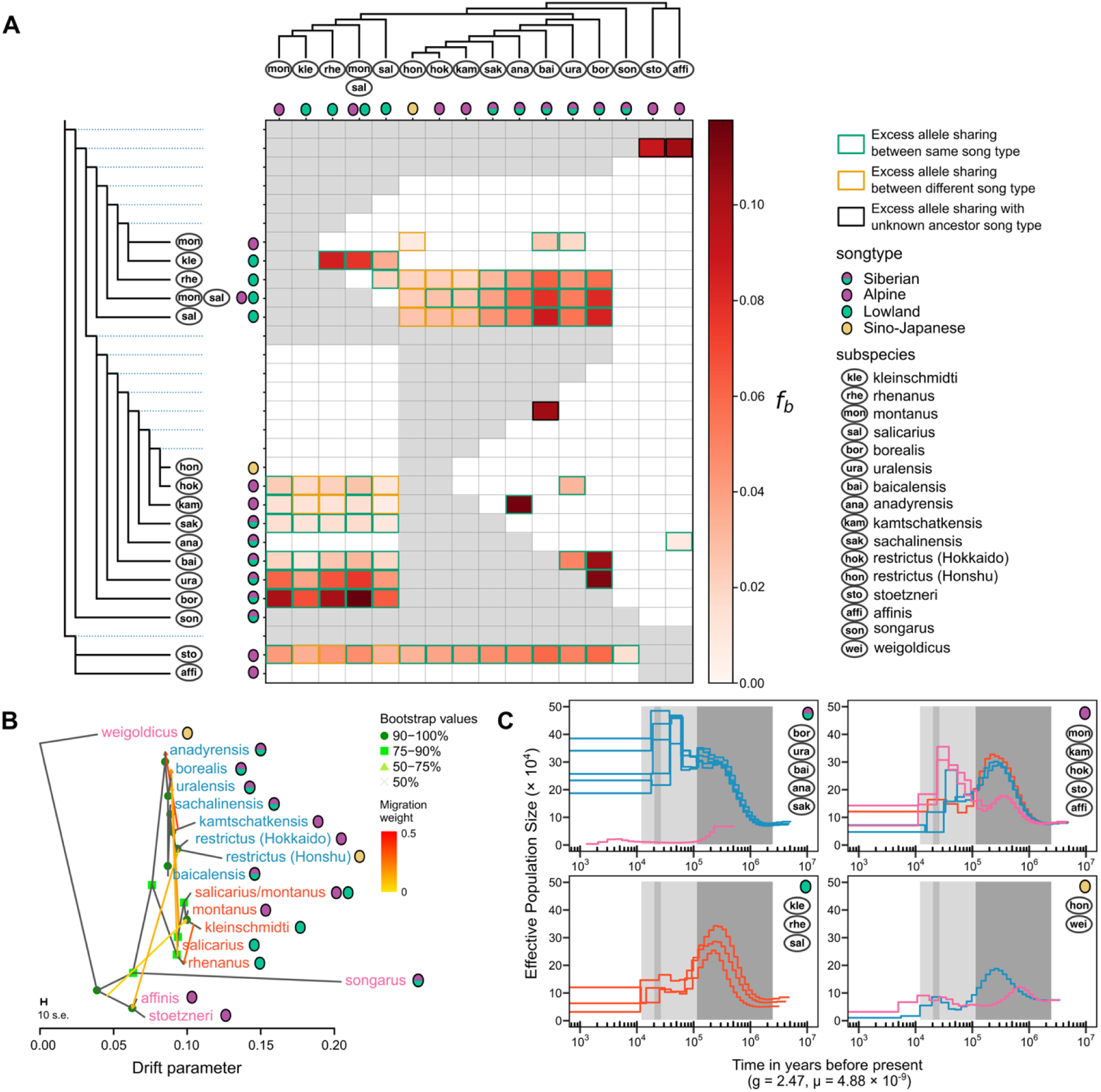
Ancestral admixture, genetic drift, and demographic history of willow tit lineages. (**A**) Excess derived-allele sharing visualised as *f*-branch statistics (ƒb(C)) among all willow tit subspecies based on 546,398 autosomal biallelic SNPs. Each cell quantifies excess allele sharing between lineage *C* (x-axis) and branch *b* (y-axis) relative to its sister branch. Cyan outline indicates excess sharing between lineages with the same song type, yellow between different song types, and black where the ancestral song type is unknown. Song type symbols correspond to those in Fig. 1. (**B**) Population splits and migration events inferred with TreeMix from the same SNP dataset. Horizontal branch lengths represent genetic drift, and coloured arrows denote migration edges; colour intensity indicates migration weight. Symbols denote bootstrap support (50–100 %). (**C**) Historical effective population size (*N*J) trajectories reconstructed for one individual per subspecies using the Pairwise Sequentially Markovian Coalescent (PSMC). Time is shown on a log_10_ scale. Shaded background intervals mark the Last Glacial Maximum (dark grey, 26–20 kya), the Last Glacial Period (medium grey, 115–11.7 kya), and the broader Pleistocene (light grey, 2.58 Mya–11.7 kya). A mutation rate of μ = 4.88 × 10^-9^ per site per generation and a generation time of 2.47 years were used for scaling.

Modelling up to six migration events (m = 1–6) revealed a stable overall topology that was consistent with the autosomal phylogeny (**Supplementary Fig. S5**). All inferred migration edges connected populations that currently share at least one song type (**Fig. 3B**). Within the Central European cluster, gene flow occurred among *kleinschmidti*, *rhenanus*, and *salicarius*—populations that share the Lowland song type—and between *salicarius* and *montanus*. In the Northern Palearctic cluster, we detected migration between Siberian singers (*anadyrensis*) and Alpine singers (*kamtschatkensis*). Across clusters, the main migration signals involved populations sharing the Lowland song type, such as *salicarius* → *borealis* and *rhenanus* → *uralensis*, consistent with limited gene flow after secondary contact as inferred earlier.

To test whether, in addition to isolation-by-distance, song-type similarity contributes to gene flow among populations, we fitted multiple regression models on distance matrices (MRM) using three complementary measures of genomic differentiation: F_ST_, absolute divergence (D_XY_), and the covariance matrix estimated by TreeMix. Song similarity was encoded as an ordinal index reflecting increasing song divergence (0 = complete song sharing, 1 = partial song sharing, 2 = no song sharing), and geographic distance was included to account for isolation-by-distance.

Song-type differences were associated with elevated F_ST_ after controlling for geographic distance (β_song_ = 0.284, 95% CI [0.152,0.416], p = 0.05; **Supplementary Table S2**), indicating reduced contemporary gene flow between populations that differ in song. In contrast, song divergence was not a significant predictor of either D_XY_ (β_song_= 0.103, 95% CI [-0.028, 0.234], p = 0.35) or TreeMix covariance (β_song_ = −0.077, 95% CI [-0.182, 0.027], p = 0.213).

This contrast across metrics suggests that song primarily influences recent or ongoing gene flow, rather than having driven the deepest lineage splits within the complex. Consistently, both D_XY_ and TreeMix covariance were more strongly predicted by geographic distance than by song (p = 0.0001 and 0.0001, respectively), and these models explained substantially more variance (R^2^ = 0.178 and 0.477) than the F_ST_-based model (R^2^ = 0.166), reflecting their sensitivity to long-term divergence and shared demographic history rather than contemporary reproductive barriers. Consistent with the MRM results, excess allele sharing according to the f-branch statistic was enriched among lineages sharing song types (Fisher’s exact test, p = 5 × 10^-4^; **Fig. 3A**).

No extant subspecies showed excess derived-allele sharing uniquely with *songarus*, suggesting that its admixed ancestry does not reflect recent hybridization. Taken together, these analyses indicate gene flow from deep ancestral lineages into peripheral populations and localised admixture among geographically proximate taxa during historical range expansions. Lineages that share the same or similar song types were more likely to exchange genes, even after accounting for geography, supporting the idea that song contributes to assortative mating and may have helped maintain lineage cohesion through time.

### Population demographic history and hindcasting

We next asked whether the isolation of peripheral, song-monotypic populations was linked to specific demographic events such as bottlenecks or declines. To infer the temporal scale of these processes, we reconstructed population history using the pairwise sequentially Markovian coalescent (PSMC) method on one high-coverage diploid genome per subspecies (**Fig. 3C**; **Supplementary Fig. S6**). In parallel, we used species distribution modelling (SDM) and hindcasting to identify potential glacial refugia and assess historical habitat availability (**Fig. 4**).

**Fig. 4.**
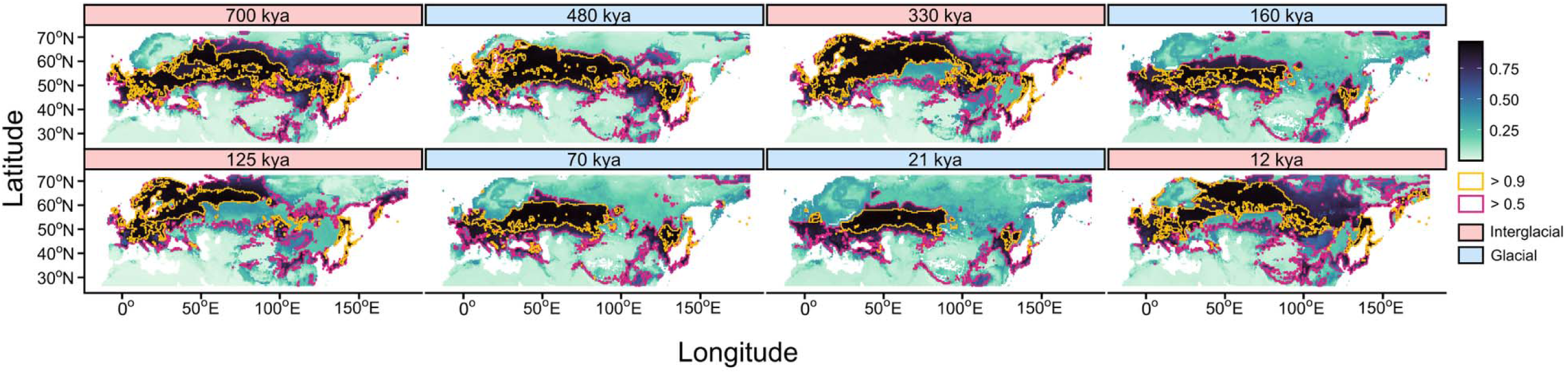
Hindcasted habitat suitability of the willow tit across representative glacial and interglacial stages during the Mid and Late Pleistocene. Species distribution models were projected onto paleoclimatic reconstructions at eight representative time slices spanning the last 0.8 Mya. Maps illustrate alternating phases of interglacial range expansion (e.g. 700, 330, 125 kya) and glacial contraction (e.g. 480, 160, 70, 21 kya), followed by post-glacial recovery after 12 kya. Colours represent modelled habitat suitability (0–1; darker colours = higher suitability). The magenta outline represents the area with >0.5 suitability, and the yellow outline shows the area with >0.9 suitability to visualise the fragmentation of suitable habitat over time. The reconstructions reveal repeated fragmentation of suitable habitat during glacial periods and extensive connectivity during interglacials, consistent with demographic patterns inferred from genomic data. Notably, regions corresponding to present-day song-monotypic populations (e.g. European Alps, Japan, Kamchatka) repeatedly experienced isolation during glacial phases, whereas the northern Palearctic retained extensive connectivity.

The earliest coalescent signals in the PSMC trajectories were observed in the southeastern Palearctic lineages—*weigoldicus*, *affinis*, and *stoetzneri*—which began to diverge during the Late Pliocene to Early Pleistocene (3.6–1.8 Mya; **Fig. 3C**). Their deep demographic histories are consistent with the phylogenetic position of these lineages as the most basal in the species tree (**Figs. 2A, 3B**) and their estimated split from the ancestors of the Northern Palearctic and Central European willow tits around 2.33 Mya (95% CI [1.73, 3.13], **Supplementary Fig. S2**), providing further support for an Asian origin of the willow tit complex. This timing coincides with known southward shifts of boreal and temperate taxa driven by early Quaternary cooling (Hewitt, 2000, 2004), and aligns with hypotheses that climatic fluctuations at the Pliocene-Pleistocene transition promoted extinction and speciation events (Avise & Walker, 1998; Lovette, 2005; Saitoh et al., 2010; Weir & Schluter, 2004).

Across all populations, effective population size increased gradually from approximately 1–0.9 Mya, coinciding with the Mid-Pleistocene Revolution (MPR), when glacial cycles shifted from 41 kya to 100 kya periodicity (Augustin et al., 2004). A comparatively stable climate between 0.9 and 0.4 Mya—with longer glacial periods and comparatively brief interglacials—likely facilitated expansion into newly available northern habitats and subsequent dispersal across the western and eastern Palearctic (**Fig. 4**; note the northward expansion of suitable habitat during interglacials).

Following this major Mid-Pleistocene expansion, ancestors of peripheral populations that now sing only one song type were likely isolated in climatic refugia, where they lost one of the ancestral song types. In Europe, Alpine and Lowland singers in the Alps, Central Europe, and Great Britain experienced marked population declines around 0.4–0.2 Mya (**Fig. 3C**). This period corresponds to the Saale and Riss glaciations (0.36–0.13 Mya) (Lauer & Weiss, 2018), when extensive ice coverage reached as far south as central Germany, plausibly fragmenting *montanus*, *rhenanus*, *salicarius*, and *kleinschmidti* populations (**Fig. 4**, note the fragmentation of suitable habitat across Europe during glacial phases).

In East Asia, rising sea levels around 0.43 Mya flooded the Tsugaru Strait, separating Honshu from Hokkaido (Shiba, 2021), while Hokkaido and Sakhalin remained intermittently connected to the mainland (Dobson, 1994; McKay, 2012). This restricted connectivity likely contributed to the genetic and song-type divergence between *restrictus* populations, consistent with phylogeographic patterns in other Japanese taxa (Honda et al., 2019; Ikeda, 2022; Sawaura et al., 2018). Similarly, isolation in Kamchatka during late Pleistocene cooling events may explain its monotypic Alpine song type (Barr & Solomina, 2014; Zink et al., 2008). Together, these patterns suggest that peripheral populations on islands (Great Britain, Honshu, Hokkaido, Sakhalin) and in mountain refugia (Alps, Kamchatka) experienced strong isolation and repeated contractions.

In contrast, the widespread Siberian singers persisted across northern Eurasia even during glacial advances (∼0.09–0.05 Mya) and maintained large effective population sizes throughout most of the Mid to Late Pleistocene. Their PSMC trajectories indicate continued growth until the onset of the Last Glacial Period (∼0.12 Mya), brief stagnation, and renewed expansion near 0.06 Mya. Climatic stability between 0.7 and 0.1 Mya likely sustained broad mid-latitude refugia (**Fig. 4**; note the persistence of contiguous suitable habitat in the north through glacial cycles), facilitating persistence and gene flow among northern subspecies, which now share the broad Siberian song repertoire.

### Implications for the role of song in lineage divergence

The extent to which willow tit song is learned remains incompletely understood. Experiments with hand-raised birds suggest that the Alpine song type may be largely innate, although these studies were not standardised and should be interpreted with caution (Thönen, 1962). At the same time, distinct song types appear to have remained stable over long evolutionary timescales. The closely related Caspian tit (*P. hyrcanus*) that sings Lowland song (Harrap & Quinn, 1995) were estimated to share a common ancestor with the willow tit around 1.38 Mya (Johansson et al., 2018). Furthermore, the Asian clade (Sino-Japanese and Alpine song) and the clade which contains Northern Palearctic and Central European willow tits were estimated to share a common ancestor as early as 2.33 Mya. More broadly, comparative evidence suggests that passerine vocalizations, including aspects of song syntax, can show strong phylogenetic signal (Arato & Fitch, 2021; Päckert et al., 2003; Tietze et al., 2015).

Despite the uncertainty in learning mechanisms, our genomic analyses show that gene flow occurred preferentially among lineages sharing similar song types, whereas lineages with distinct songs remained largely isolated. This pattern is consistent with song acting as a partial barrier to gene flow and contributing to the maintenance of lineage boundaries.

Demographic history further modulates this relationship. Peripheral populations that experienced repeated bottlenecks during glacial isolation are associated with monotypic repertoires, consistent with cultural drift in small, isolated populations (Baker et al., 2001; Ciaran White, 2012; Crates et al., 2025; Koetz et al., 2007; Lachlan et al., 2013; Lynch, 1996; Paxton et al., 2019). In contrast, large and well-connected northern populations retained broad mixed (‘Siberian’) repertoires, reflecting long-term connectivity and cultural exchange.

## Conclusions

Our analyses indicate that the willow tit complex originated in Asia during the Late Pliocene to Early Pleistocene and expanded across the Palearctic during subsequent interglacial periods. This scenario helps reconcile earlier contrasting hypotheses of westward (Kvist et al., 2001) versus eastward (Pavlova et al., 2006) postglacial expansion, which likely arose from incomplete geographic sampling. By combining genome-wide data with full-range coverage, we provide a unified view of expansion from Asia followed by dispersal across the Palearctic. This history resulted in three deeply diverged genomic lineages—Asian, Central European, and Northern Palearctic—whose boundaries coincide with major song-type divisions. Historical climatic shifts likely created alternating barriers and corridors to gene flow across the Palearctic. Hindcasting indicates that suitable habitat remained extensive in northern regions but fragmented at the range periphery. This pattern suggests that periods of connectivity maintained broad mixed (‘Siberian’) song repertoires in the north, whereas repeated isolation in refugia was associated with song monotypy at range margins, as seen on Honshu, Hokkaido, Great Britain, and the European Alps, which were likely more isolated and may have diverged further through genetic drift, while the more widespread northern Palearctic subspecies remained connected by gene flow.

Taken together, these findings demonstrate that genomic divergence and learned song have been tightly linked throughout the evolutionary history of the willow tit. Learned song, while culturally transmitted, has mirrored genetic lineage boundaries and likely contributed to their maintenance over hundreds of thousands of years, providing evidence for gene-culture coupling in a widespread passerine.

## Materials and Methods

### Samples and song type assignment

For the reference genome, blood was sampled from two female willow tits (*Poecile montanus salicarius*) captured in southern Germany (48.10° N, 11.35° E) in 2021. For population resequencing, tissue, blood, or footpad samples were obtained from museums and individual researchers. Subspecies and song type were assigned from sampling localities (Table 1) using the range synthesis in Quaisser & Eck (2002), which compiles the most up-to-date distributions of subspecies and song types. Samples that were located on a few areas where pure Alpine/Lowland or Sino-Japanese singers locally co-occur, i.e. in the Altai mountains, or in the Czech Republic near the border with Austria and Germany (Fijen, 2015; Martens et al., 1995; Thönen, 1996), were assigned ambiguous song type (i.e. “Alpine/Lowland”).

For the population genomic analyses, we analysed 88 willow tit samples covering the entire distribution range and all fourteen recognised subspecies: *songarus* (*N* = 2), *stoetzneri* (*N* = 7), *affinis* (*N* = 9), *restrictus* - Honshu (*N* = 4), *restrictus* - Hokkaido (*N* = 3), *sachalinensis* (*N* = 4), *anadyrensis* (*N* = 5), *kamtschatkensis* (*N* = 4), *baicalensis* (*N* = 12), *uralensis* (*N* = 5), *borealis* (*N* = 10), *salicarius* (*N* = 2), *salicarius/montanus* (*N* = 7), *montanus* (*N* = 5), *rhenanus* (*N* = 6), and *kleinschmidti* (*N* = 3, **Supplementary Table S3**). We additionally included three Sichuan tit (*P. weigoldicus*) individuals and two marsh tit (*P. palustris*) samples that were used as an outgroup in the phylogenetic analyses.

### Reference genome assembly and annotation

#### DNA extraction and sequencing

High-molecular-weight DNA for PacBio HiFi sequencing was extracted with the Monarch HMW DNA Extraction Kit for Cells & Blood (New England Biolabs). Illumina short-read libraries were prepared from the same individual. For Hi-C, DNA was extracted from fresh-frozen blood of a second female and processed with the Arima Genomics Hi-C kit (two technical replicates). We generated 58.1 Gb PacBio HiFi data (∼47.5× coverage for a 1.2 Gb genome), 118.9 Gb Hi-C data in total (∼100× coverage), and 64.9 Gb Illumina short-read data (∼54× coverage).

#### Assembly, scaffolding, and curation

HiFi reads were assembled with *hifiasm* v0.16.1-r375 (Cheng et al., 2021) in *primary* mode using default settings. Initial assembly contiguity, completeness, and accuracy were evaluated with *QUAST* v5.1.0rc1 (Gurevich et al., 2013) and *BUSCO* v5.2.2 (Manni et al., 2021; Simão et al., 2015) in *genome* mode using the aves_odb10 database, and base-level quality with *meryl* v1.3 with 21 bp as the k-mer size (Rhie et al., 2020). Hi-C reads were trimmed and quality-checked with *fastp* v0.23.2 (Chen et al., 2018), *qc3c* v0.5 (DeMaere & Darling, 2021), and *MultiQC* v1.10 (Ewels et al., 2016). Following the *Arima Hi-C mapping pipeline A160156 v02* (Arima Genomics, 2019), reads were mapped with *BWA MEM* v0.7.17-r1188 (Li & Durbin, 2009), and processed to remove chimeric reads and PCR duplicates using *Picard v2.27.5* (Broad Institute, 2019). Libraries were merged using *Picard MergeSamFiles*. Scaffolding used *yaHS* v1.1a-r3 (Zhou et al., 2023) with default settings, followed by manual mis-join correction in *Juicer* v1.1 and *Juicebox Assembly Tools* v1.11.08 (Dudchenko et al., 2017, 2018). Putative sex chromosomes were assigned based on Hi-C read coverage and homology to other avian genomes and were removed from further analyses. Using *RagTag v2.1.0* (Alonge et al., 2019, 2022), scaffolds were ordered/oriented to the great tit (NCBI Accession no. GCF_001522545.3) and zebra finch (NCBI Accession no. GCA_003957565.4) assemblies to form chromosome-scale pseudomolecules. Final genome assembly metrics (**Supplementary Fig. S7, Supplementary Table S4**) were visualised using *blobtoolkit v3.2.7* (Laetsch & Blaxter, 2017).

#### Repeat annotation and masking

Interspersed repeats were identified with *RepeatModeler2* v2.0.5 (option ‘-LTR_struct’). Following the protocol typically used for avian genomes, the resulting de novo repeat library was merged with *Dfam v3.8* (Storer et al., 2021), *RepBase v20181026* (Bao et al., 2015), and published passerine repeat libraries (blue-capped cordon-bleu (Boman et al., 2019), collared flycatcher (Suh et al., 2018), hooded crow (Weissensteiner et al., 2020), Anna’s hummingbird, emu, kākāpō (Peona, Palacios-Gimenez, et al., 2021), paradise crow (Peona, Blom, et al., 2021), and eastern black-eared wheatear (Peona et al., 2023)). The combined library was used to soft-mask the assembly with *RepeatMasker v4.1.6* using ‘-xsmall and -gff’ parameters (Smit & Hubley, 2023).

#### Mitochondrial genome

The mitochondrial genome was assembled with *MitoFinder* v1.4.1 (Allio et al., 2020) under default settings using a pre-existing willow tit mitochondrial reference genome (NCBI GenBank accession no. MN122849). Mitochondrial genes were annotated with *MitoZ* v3.4 using the ‘--clade Chordata’ parameter (Meng et al., 2019).

### Population genomic resequencing

#### DNA extraction and sequencing

Genomic DNA from all 93 samples was extracted using the GeneON GF.1 Blood Extraction Kit (Vivantis) and the Monarch Genomic DNA Purification Kit (New England Biolabs) following the manufacturer’s protocols. Library preparation and sequencing were outsourced and samples were sequenced to an average depth of approximately 20 Gb (∼17 × coverage; mean ± SD = 17.56 ± 3.86) using the Illumina NovaSeq platform (150 bp paired-end reads with 400 bp insert size).

#### Read processing and variant calling

All resequencing data were run through *grenepipe* v0.12.2 (Czech & Exposito-Alonso, 2022) using the default settings if not otherwise stated. Sequences from each sample library were quality-checked with *FastQC* v0.11.9 (Andrews, 2020), and subsequently trimmed with *cutadapt* v3.5 (Martin, 2011) by providing the adapter sequences and a quality cut-off for the 5’ and 3’ ends before adapter removal. Then, reads were mapped to the willow tit reference genome using *BWA-MEM* (Li & Durbin, 2009) using the option ‘-R’ to use the read groups tags, as well as ‘-M’ for compatibility with Picard. Duplicates were marked with *Picard MarkDuplicates* (Broad Institute, 2019). Alignment quality was assessed with *samtools* v1.12 (*stats*, *flagstat*) (Danecek et al., 2021), *Qualimap* v2.2.2a (García-Alcalde et al., 2012; Okonechnikov et al., 2016), and *Picard CollectMultipleMetrics* (Broad Institute, 2019).

Variants were called using *GATK* v4.1.4.1 *HaplotypeCaller* in gVCF mode and were then genotyped using *GenotypeGVCFs* (DePristo et al., 2011; McKenna et al., 2010). Hard-filtering parameters were applied per GATK recommendation separately for SNPs and Indels (Van der Auwera et al., 2013). For SNPs, we filtered with the following filtering parameters: QD < 2.0, FS > 60.0, MQ < 40.0, MQRankSum < −12.5, ReadPosRankSum < −8.0; for indels: QD < 2.0, FS > 200.0, ReadPosRankSum < −20.0. For invariant sites, we applied the following filter: QUAL < 15 || DP < 150.0, based on invariant site quality and across-samples read depth. Hard-filtered sites and sites overlapping repeats were removed from the dataset prior to further filtering.

#### Variant filtering and masking

From the initial hard-filtered dataset (59,292,222 SNPs and indels), biallelic autosomal SNPs were retained (48,761,502 SNPs) and genotypes with GQ < 15 or read depth < 5 were set to missing (“.”). Subsequently, sites with >20% missing data or minor allele frequency < 0.05 were removed with *VCFtools* v0.1.16 (Danecek et al., 2011), yielding 10,359,256 biallelic SNPs (see below for the exact subsets used per analysis, as some required further pruning based on linkage disequilibrium).

### Mitochondrial phylogeny reconstruction

Whole mitochondrial genomes of all 91 individuals (88 *P. montanus* and 3 *P. weigoldicus*) and the outgroup *P. palustris* (NCBI GenBank accession no. <u>NC_026911.1</u>) were aligned using *MAFFT* v6.864b with 1,000 iterative refinements (Katoh & Standley, 2013). The best-fitting nucleotide substitution model was identified using *ModelTest-NG* v0.1.7 (Darriba et al., 2020), which selected the general time-reversible model with invariant sites and gamma-distributed rate variation (GTR + I + G4). A maximum-likelihood (ML) phylogeny was inferred with *RAxML-NG* v1.2.1 (Kozlov et al., 2019; Stamatakis, 2014) under the GTR + I + G4 model with 1,000 bootstrap replicates.

A Bayesian phylogeny was inferred with *MrBayes* v3.2.7a (Ronquist et al., 2012) using two independent runs of four Markov chains (three heated, one cold) for 10,000,000 generations, sampling every 1,000^th^ generation. The first 25% of samples were discarded as burn-in, and convergence was confirmed by an average standard deviation of split frequencies < 0.01 and effective sample sizes > 200 using *Tracer* v1.7.2 (Rambaut et al., 2018).

Phylogenetic trees were visualised in *R* v4.5.1 using the packages *treeio* v1.32.2, *ggtree* v3.16.3, and *ggplot2* v4.0.1 (R Core Team, 2025; Wang et al., 2020; Wickham, 2016; Yu et al., 2017). For both the maximum likelihood and Bayesian phylogeny, the outgroup was used to root the phylogeny, and subsequently dropped for simplification and reduced tree length.

### Autosomal phylogeny and divergence time estimation

Two *P. palustris* samples were included in the autosomal phylogeny construction as an outgroup. For constructing the gene trees, autosomal variants were pruned for linkage disequilibrium (r^2^ > 0.1) within 50-SNP-windows and a 10-SNP-step using *PLINK* v1.90b6.21 (Chang et al., 2015) with the parameters ‘--indep-pairwise 50 10 0.1’, resulting in 1,374,753 SNPs. The variants were haploidised using a modified version of ‘<u>phyml_sliding_windows.py</u>’ (Jensen et al., 2023; Martin, 2020) by selecting a random allele at heterozygous sites. Gene trees were constructed from autosomal chromosomes using *IQ-TREE* v2 (Minh et al., 2020) for 50kb windows sampled every 1Mb with 1000 ultrafast bootstrap support, which resulted in 1,869 gene trees. The General Time Reversible (GTR) model of sequence evolution was chosen with an ascertainment bias correction to account for the missing invariant sites. A coalescence-based species tree was inferred from the resulting gene trees using *ASTRAL* (Mirarab & Warnow, 2015) with 25 rounds of placement and subsampling for each exploration step.

Divergence times were estimated using an approximate likelihood method implemented in *MCMCTree* v4.10.8.iq2mc (Reis & Yang, 2011). Calibration information from Johansson et al. (2018) was added at the split between *P. palustris* and the willow tit on the *ASTRAL* topology with a soft minimum and maximum bound (2.5% probability of exceeding the bounds). To lessen computational burden, 20 50kb autosomal windows were randomly selected and aligned using *MAFFT*. The *IQ2MC* workflow (Demotte et al., 2025) as implemented in *IQ-TREE* v3 (Wong et al., 2025) was used to estimate the gradient and Hessian of the branch lengths of the phylogeny using the GTR substitution model. Four independent runs of Markov chains were run for 10,000,000 generations, sampled every 1,000^th^ generation with the first 1,000,000 generations discarded as burn-in. Convergence was checked using *Tracer* by ensuring that effective sample size was > 200 and that acceptance proportions in the output file were between 20–40%.

### Autosomal population structure: Principal component and admixture analysis

To complement the mitochondrial phylogeny and assess genome-wide population structure, autosomal SNPs from the linkage pruned variants dataset were used. Principal component analysis (PCA) and ancestry inference were first conducted including all 91 individuals (88 *P. montanus* and 3 *P. weigoldicus*) to capture overall genetic structure across the species complex. A second PCA excluding the Asian lineages (*weigoldicus*, *affinis*, *stoetzneri*, *songarus*) was performed to resolve finer-scale differentiation among the Central European and Northern Palearctic populations (*N* = 72 individuals).

PCA was carried out in *PLINK*, and ancestry proportions were estimated in *ADMIXTURE* v1.3.0 (Alexander et al., 2009). Models with K = 1–10 ancestry clusters were tested, with each analysis repeated ten times using different random seeds. PCA results were visualised in *R* using *ggplot2*, and *ADMIXTURE* results were summarised with *pong* v1.5 and spatially interpolated with *TESS3* v1.1.0 (Behr et al., 2016; Caye et al., 2016; Wickham, 2016).

### Historical gene flow: migration and introgression tests

Analyses of historical population splits and migration were based on the filtered and LD-pruned autosomal dataset after removing sites with missing data, resulting in 546,398 SNPs. We pooled samples based on subspecies designation, which was based on geographic sampling location and independent of song type. This allowed testing whether song type has any effect on gene flow.

Population splits and migration edges were inferred using *TreeMix* v1.13 with the ‘-se’ parameter for standard error calculation (Pickrell & Pritchard, 2012). Covariance matrices were estimated in blocks of 10,000 SNPs to account for linkage disequilibrium. A consensus tree was built from 100 bootstrap tree replicates and rooted with *P. weigoldicus*. Models including 0–10 migration edges were evaluated, each replicated ten times, and the optimal range (m = 0–3) was identified using *OptM* (Fitak, 2021). To allow for the complex geographic history of the species, final analyses allowed up to six migration edges (m = 0–6), each with 30 replicates. For each value of m, the tree with the highest likelihood was retained and visualised using custom *R* scripts incorporated in *TreeMix_functions.R* (Dahms, 2021), adopted from Zecca et al. (2020). Examination of the residual covariance matrix (**Supplementary Fig. S8**) indicated that the model with six migration edges provided an adequate fit to the observed allele-frequency covariance. Most population pairs were well explained by the inferred topology, and residuals were generally small, with only a few localised deviations exceeding ±3 standard errors. These residuals occurred primarily among geographically proximate populations, consistent with fine-scale structure or weak additional covariance not explicitly modelled by TreeMix.

Introgression was tested between all possible lineage triplets using Patterson’s D-statistics (ABBA-BABA) (Patterson et al., 2012) and the related f_4_-ratio statistic implemented in *Dsuite* v0.5 r57 (Koppetsch et al., 2024; Malinsky et al., 2021), on the TreeMix consensus topology with *weigoldicus* as the fixed outgroup. Excess allele sharing was visualised with the f-branch statistic ƒb(C) (Malinsky et al., 2018, 2021), which quantifies derived-allele sharing between lineage *C* (x-axis) and branch *b* relative to its sister branch (y-axis). Bonferroni-corrected *p*-values were computed in the *stats* package in *R* (R Core Team, 2025).

Migration patterns were visualised under a spatially heterogeneous isolation-by-distance model using Fast Estimation of Effective Migration Surfaces (FEEMS) (Marcus et al., 2021; Petkova et al., 2015). This method is particularly useful for revealing population structure that deviates from a pure isolation-by-distance model, such as an extension of barrier structures, or evolutionary processes, such as natural selection or drift. The migration surface was extrapolated on a triangular lattice comprised of nodes (‘subpopulation’) and weighted edges with a cell size of 25,000 km^2^ and a cell spacing of 220 km. The edges were weighted based on estimated symmetric gene flow between ‘subpopulations’. The smoothing parameters used to penalise differences between the estimated edges were determined using a 5-fold cross-validation procedure for all pairwise combinations of 10 and 5 different values of λ and λ_q_ (**Supplementary Fig. S4**).

### Population genomic summary statistics

Population genetic diversity and differentiation were quantified using the repeat-masked and filtered variants along with the hard-filtered invariant sites. Nucleotide diversity (π) was calculated for each subspecies, and genetic differentiation (F_ST_), and absolute divergence (D_XY_) were calculated for each subspecies pair (**Supplementary Fig. S9–S10**) using *popgenWindows.py* (Martin, 2020; Martin et al., 2019; Merrill et al., 2019). Estimates were computed in non-overlapping 50 kb windows containing a minimum of 30 variable sites using the formula for K_ST_ (Hudson et al., 1992) because it is less sensitive to small sample size.

### Association between song type and gene flow

The association between song-type similarity and genomic differentiation was assessed using multiple regression on distance matrices (MRM), implemented in the *algatr* package (Chambers et al., 2025; Wang, 2013). Pairwise genetic differentiation (F_ST_) was used as the response variable. Song-type similarity was encoded as an ordinal distance matrix: 0 = complete song sharing, 1 = partial song sharing (e.g. among mixed Siberian singers and both Alpine and/or Lowland singers), 2 = no song sharing (e.g. among Sino-Japanese singers and singers displaying any other song type). In all analysis using song type, samples in Central Europe that were found in regions where both Alpine and Lowland song co-exist, are treated the same as Siberian singers but denoted as Alpine and Lowland song type (**Fig. 2A, 3A–B, Supplementary Fig. S1, S2**). Geographic distance (log-transformed great-circle distance between population centroids) was included as a covariate to account for isolation-by-distance. Equivalent models were fitted using absolute divergence (D_XY_) and the allele-frequency covariance matrix estimated by TreeMix as alternative response variables. Statistical significance was assessed using 10,000 permutations. Relationships between variables were visualised using the *mmrr_plot* function (**Supplementary Fig. S11**).

To assess whether excess allele sharing inferred from f-branch statistics was non-randomly associated with song-type similarity, f-branch values were categorised according to whether the corresponding population pairs shared the same (*N* = 94), partial (*N* = 154), or no (*N* = 75) song types. Internal branches were not assigned song types, and all f-branch matrix entries were treated as potential allele-sharing events (*N* = 510). A contingency analysis was used to test for enrichment of excess allele sharing among song-sharing lineages using Fisher’s exact test in *R* (R Core Team, 2025). Because f-branch entries are not statistically independent, this analysis was interpreted descriptively rather than as a formal test.

### Demographic history reconstruction

Demographic history was reconstructed for one individual per subspecies using the Pairwise Sequentially Markovian Coalescent (*PSMC* v0.6.5-r67) model (Li & Durbin, 2011). Consensus genome sequences were generated from mapped reads with *bcftools* v1.15 (Danecek et al., 2021) using the *mpileup* function, retaining sites with read depth between 10 and 100 and a minimum root-mean-squared mapping quality of 20. *PSMC* analyses were run for 25 iterations (-N 25), 10 generations back in time (-t 10), and an initial θ/ρ ratio of 5. The parameter pattern was set to ‘-p 2+2+25*2+4+6’, defining 29 free parameters across 64 atomic intervals. This configuration splits the first-time window, as recommended by Hilgers et al. (2025), to avoid false recent population peaks arising from ill-set default parameters. Several alternative ‘p’ parameters from previous publications were also tested to confirm that inferred demographic trajectories were robust to parameter choice.

Parameters ‘-p’ and ‘-t’ were adjusted to ensure that at least 10 recombination events were inferred within each interval after 20 iterations (Li & Durbin, 2011). Time and effective population size estimates were scaled using a per-site, per-generation mutation rate of 4.88 × 10^-9^ as estimated for the closely related blue tit in Bergeron et al. (2023), and a generation time of 2.47 years for the willow tit (Bird et al., 2020). For each individual, 100 PSMC bootstrap replicates were performed, and median trajectories were used for visualization.

### Species distribution models and hindcasting

Species occurrence data were compiled from public databases, including Xeno-Canto, Macaulay Library, the Global Biodiversity Information Facility (GBIF), and Integrated Digitised Biocollections (iDigBio), using the search terms ‘*Poecile montanus*’ and ‘*Parus montanus*’. Additional records were extracted from previous studies (Quaisser & Eck, 2002; Tritsch et al., 2017). After removal of duplicates and records located on water bodies, the full dataset comprised 218,687 occurrence records. Environmental predictors were obtained from the high-resolution (0.5° × 0.5°) global bioclimatic (BIOCLIM) dataset of Krapp et al. (2021), which provides 19 standard bioclimatic variables summarizing temperature and precipitation means and extremes, as well as biome-related variables, for the past 0.8 Mya at 1 kya temporal resolution. The study extent covered the entire distribution range of the willow tit (−14° to 180° longitude, 26° to 72° latitude), represented as the same 0.5° × 0.5° grid as the BIOCLIM dataset (35,696 cells, of which 23,880 corresponded to terrestrial habitat after masking).

### Predictor selection

To select a consistent set of environmental predictors for species distribution modelling, predictor collinearity was assessed once using a presence-only approach. A preliminary presence-only species distribution model was fitted using all available occurrence records (*N* = 218,687) with the maxlike() function in the *maxlike R* package (Royle et al., 2012). Based on this model, grid cells with very low predicted occurrence probability (< 5 × 10^-6^) were assumed to represent environmentally unsuitable background, from which 15% of cells were randomly sampled as pseudoabsence points (*N* = 14,218). All 19 BIOCLIM variables and additional biome-related predictors were extracted for all presence and pseudoabsence points used in this step. Pairwise correlations among continuous predictors were assessed using Pearson’s r, and variables with |r| > 0.7 were excluded using the *select07* function implemented in *mecofun* v0.7.1 (Zurell, 2020). Multicollinearity among the remaining predictors was evaluated using generalised variance inflation factors (GVIF < 4) in the *car* package (Fox & Weisberg, 2019). This procedure resulted in a final set of nine predictors: Annual Mean Temperature (BIO1), Temperature Seasonality (BIO4), Mean Temperature of the Wettest Quarter (BIO8), Precipitation of the Wettest Month (BIO13), Precipitation of the Driest Month (BIO14), Precipitation Seasonality (BIO15), global biome distribution, annual net primary productivity, and leaf area index (Kaplan et al., 2003; Krapp et al., 2021). This predictor set was retained for all subsequent model replicates.

### Species distribution modelling and hindcasting

For species distribution modelling, occurrence records were spatially thinned to one presence per grid cell, resulting in 4,067 unique presence points. For each of 30 model replicates, an equal number of pseudoabsences was randomly sampled from terrestrial grid cells located outside a 300 km buffer around presence points to account for uncertainty in pseudoabsence placement. Data were split into training (80%) and testing (20%) subsets. Models were fitted using four algorithms: random forest (Breiman, 2001), generalised linear models (*stats* R package), boosted regression trees, and maximum entropy, as implemented in the *dismo* R package v1.3-15 (Hijmans et al., 2024; Phillips et al., 2006). Model performance was evaluated using the area under the receiver operating characteristic curve (AUC-ROC) via the evalSDM function in *mecofun* R package (Zurell, 2020). Thirty replicates per algorithm were run (120 models total), and an ensemble prediction was generated as a weighted mean of all models, with weights proportional to AUC-ROC values (**Supplementary Table S5**). The final ensemble was first projected onto present-day climatic layers (0 kya; **Supplementary Fig. S12**), yielding predictions consistent with the known current distribution of the willow tit (**Fig. 1A**, **B**). The ensemble was then projected onto paleoclimatic layers back to 0.8 Mya to reconstruct historical species distributions and identify potential glacial refugia through time.

## Data availability

Reference genome for the willow tit is available on NCBI under the following accessions number: <u>GCA_038429875.2</u>, and its associated raw sequences under the following project number: <u>PRJNA985397</u>. Population resequencing dataset is available on NCBI SRA under the following project number: <u>PRJNA986184</u>.

## Code availability

Codes to reproduce the results is available via Github at https://github.com/athenasyarifa/willowtit-project.git and FigShare at https://figshare.com/s/edff0a272ea3df9a041e.

## Acknowledgments

This research was funded by the Deutsche Forschungsgemeinschaft (DFG, German Research Foundation) – project number 521246704 to U.K.. Computational analyses were performed on the BioHPC hosted at Leibniz Rechenzentrum Munich funded by the Deutsche Forschungsgemeinschaft (DFG, German Research Foundation) – grant INST 86/2050-1 FUGG to J.B.W.W.. Feldbausch Foundation at Fachbereich Biologie, Mainz University, regularly supported J.M. with grants to carry out fieldwork in Asia, mainly China.

We are grateful to Shigeki Asai and the Yamashina Institute for Ornithology for providing the *Poecile restrictus* samples, Hein Van Grouw and the Natural History Museum in Tring for providing the *P. kleinschmidti* samples, and the following persons who originally collected the samples provided to Jochen Martens, Martin Päckert and Laura Kvist: Aleksandr Nazarenko (Birakan and Ussuri River and Chaplanovo, Russia), Anna Thessing (Tovetorp, Sweden), Evgeniy Georgievich Lobkov (Kamtchatka, Russia), Indrikis Krams (Latvia), Larissa Zelenskaya (Magadan, Russia), Leonid Portenko (Markovo, Russia), Markku Orell (Oulu, Finland), M. Gottschlich (Horka, Germany), Siegfried Eck (Bressanone, Italy), Stephan Ernst (Chagan-Uzun, Russia), Stefan Ferger (Eich am Rhein, Germany), Dieter Thomas Tietze (Eich am Rhein, Germany), Thomas Hallfarth (Lake Khövsgöl, Mongolia), Vadim Feodorof (Kamensk-Uralski and Kuzino, Russia), Wang Xiaoai (Qilian Mountains, China), Willfried Hansen (Hannover, Germany), Tobias Stenzel (Shaamar, Mongolia), Zbigniew Bochenski (Samjiyon, North Korea). We thank Gaby Kumpfmüller for lab work, and Javier Lazaro for his illustrations of the Poecile subspecies.

## Author contributions

A.S. and U.K. conceptualized the study. J.M., M.P., L.K., Y.H.S., L.W. contributed critical samples and materials. A.S. analysed the data, with input from U.K. and J.B.B.W.. A.S. wrote the original draft of the manuscript. U.K. and J.B.B.W supervised the study. All authors reviewed, edited, and approved the manuscript.

