## Supplementary Figures for "Cultural transmission and genomic co-divergence in the willow tit across the Palearctic"

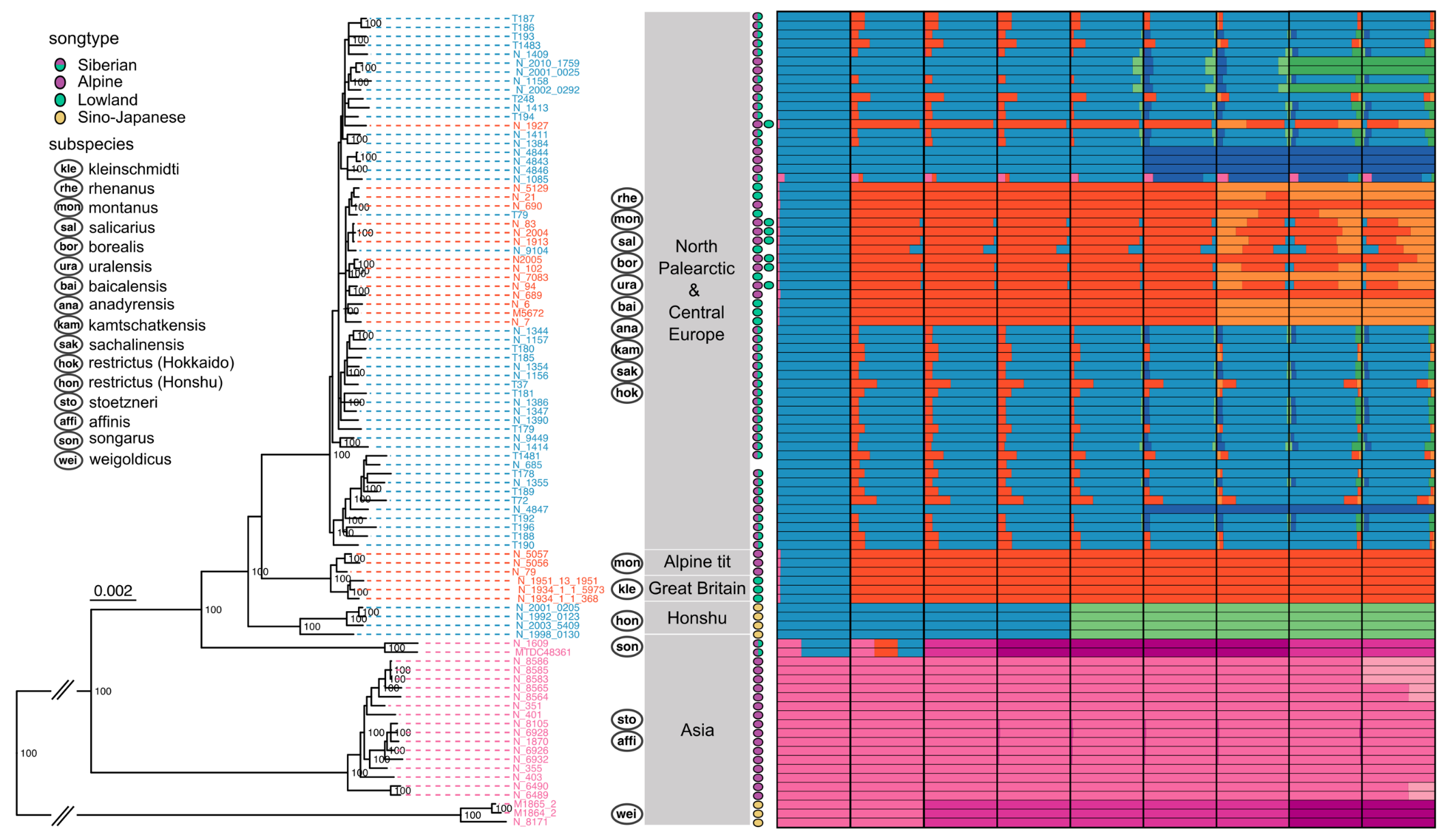

**Supplementary Figure S1.** Bayesian mitogenome phylogenetic tree inferred using MrBayes (v3.2.7a; Ronquist et al., 2012). We summarised two independent runs using the general time reversible substitution model (GTR + I + G4) with 10,000,000 steps of Markov-chain Monte-Carlo (MCMC, 25% burn-in), sampled every 1,000 generations. The phylogenetic tree was rooted with *Poecile palustris* (NCBI GenBank Accession no. NC_026911.1), although not shown on the tree for simplification and reduced tree length. Posterior probability support is shown as a percentage and is displayed only for nodes with 100% support. Geographic clusters are described on the right side, along with the subspecies that can be found within the group, followed by song type symbols for Alpine, Lowland, Siberian (Alpine + Lowland), and Sino-Japanese songs. These are followed by admixture ancestry plots from the results of ADMIXTURE v1.3.0 (Alexander et al., 2009) based on 1,374,753 autosomal SNPs. Bars represent each sample on the phylogenetic tree, and the colours represent the ancestry at K = 2–10 to the right.

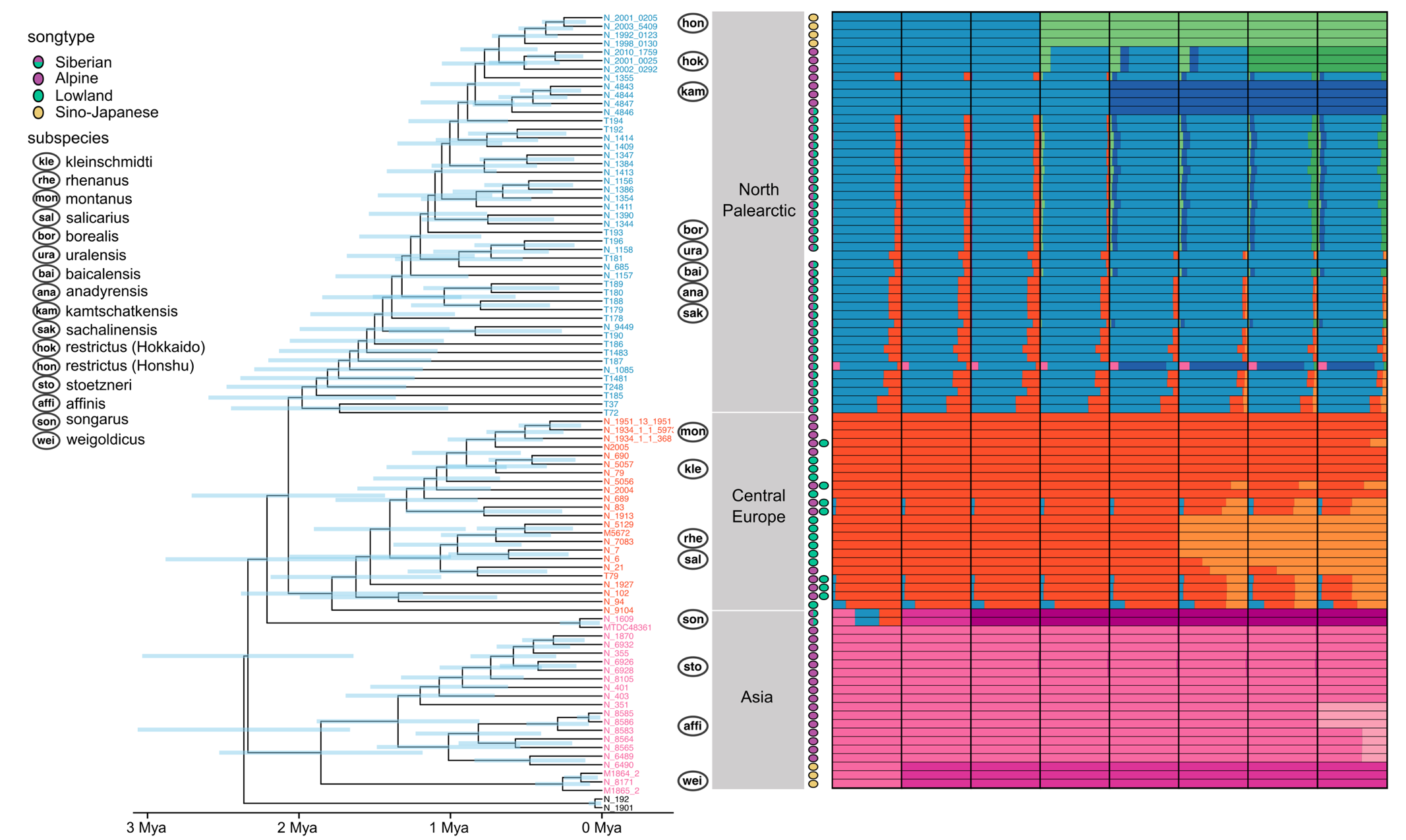

**Supplementary Figure S2.** Scaled coalescence-based species tree estimated using ASTRAL from 1,869 gene trees assembled from 50kb windows sampled every 1Mb across the whole genome using IQTREE. For heterozygous sites, a random allele was chosen for the haploidisation of the variants, and the tree support was estimated using 1,000 ultrafast bootstraps. Branch lengths are in coalescence unit and quartet branch support (fractions of quartet species trees that support the branching) above 0.5 are shown. Blue bars on the phylogeny represent the 95% CI of the divergence time estimated for the respective node by MCMCTree (Reis & Yang, 2011). On the right, subspecies for clades in the phylogeny, geographic clusters, and song types are shown. Finally, ancestries estimated for K = 3–10 are shown.

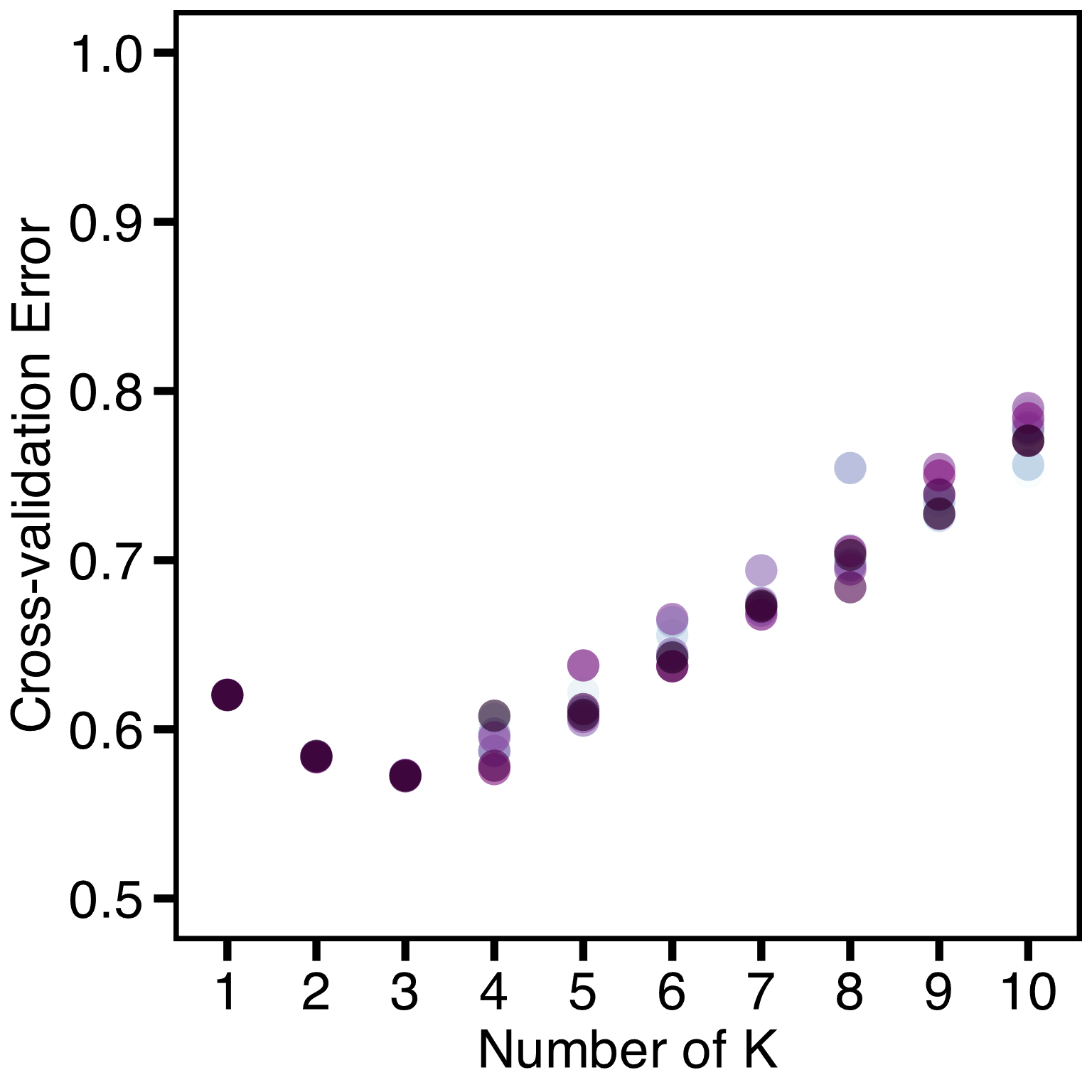

**Supplementary Figure S3.** Cross-validation standard errors for Admixture models K = 1–10. Admixture uses a 5-fold cross-validation procedure to predict the accuracy of the model, which involves partitioning the observed genotypes into 5 equally sized folds. The procedure masks genotypes of each fold in turn and uses the unmasked dataset to predict the masked genotypes. Then, by averaging the squares of the deviance residuals for the binomial model for all turns, it estimates the accuracy of the dataset for each K.

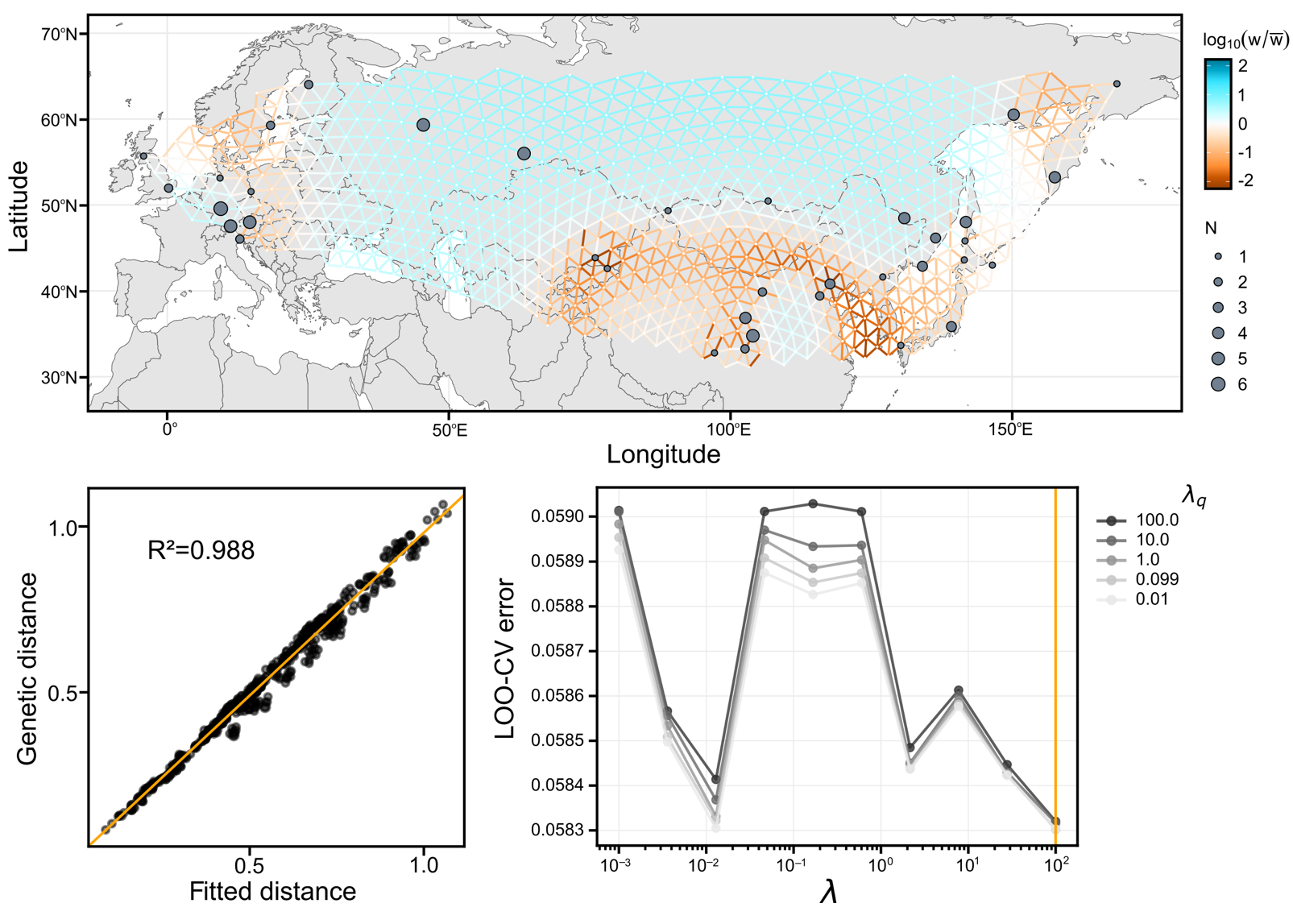

**Supplementary Figure S4.** Fast Estimation of Effective Migration Surfaces (FEEMS) (Marcus et al., 2021) using a SNP dataset without missing data (546,398 SNPs). Smoothing parameters for the model (λ = 100 and λ_q_ = 0.1) were used to penalise differences between the estimated edges on a triangular lattice comprised of nodes (‘subpopulations’) and weighted edges with a cell size of 25,000 km^2^ and a cell spacing of ~220 km.

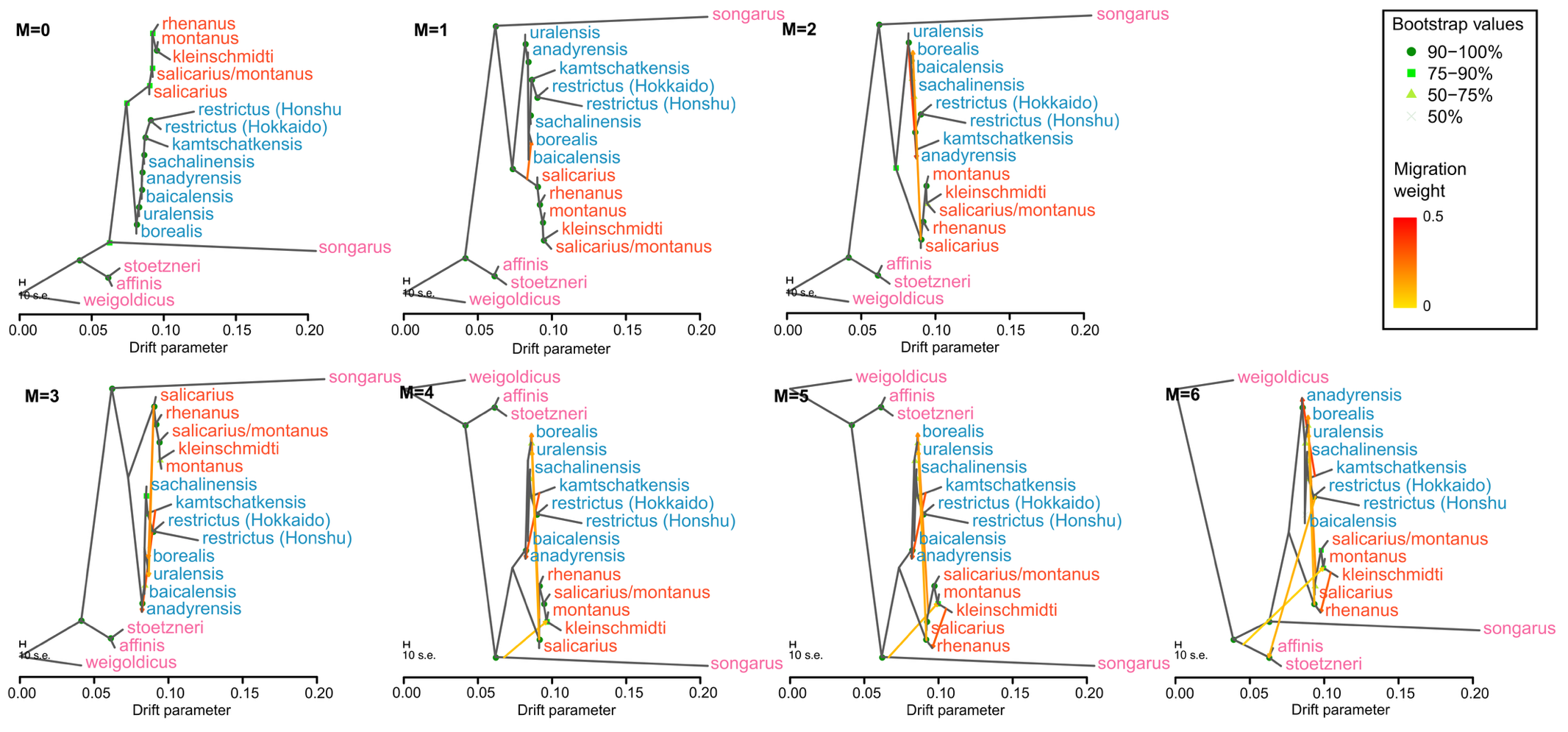

**Supplementary Figure S5.** Maximum likelihood phylogeny from TreeMix v1.13 (Pickrell & Pritchard, 2012) models with 1 to 6 migration edges and without added migration edges using 546,398 autosomal SNPs. A consensus maximum likelihood phylogeny was built from 100 bootstrap tree replicates, and subsequently migration edges were added, each with 10 replicate runs. The phylogeny with the highest likelihood was chosen for visualization.

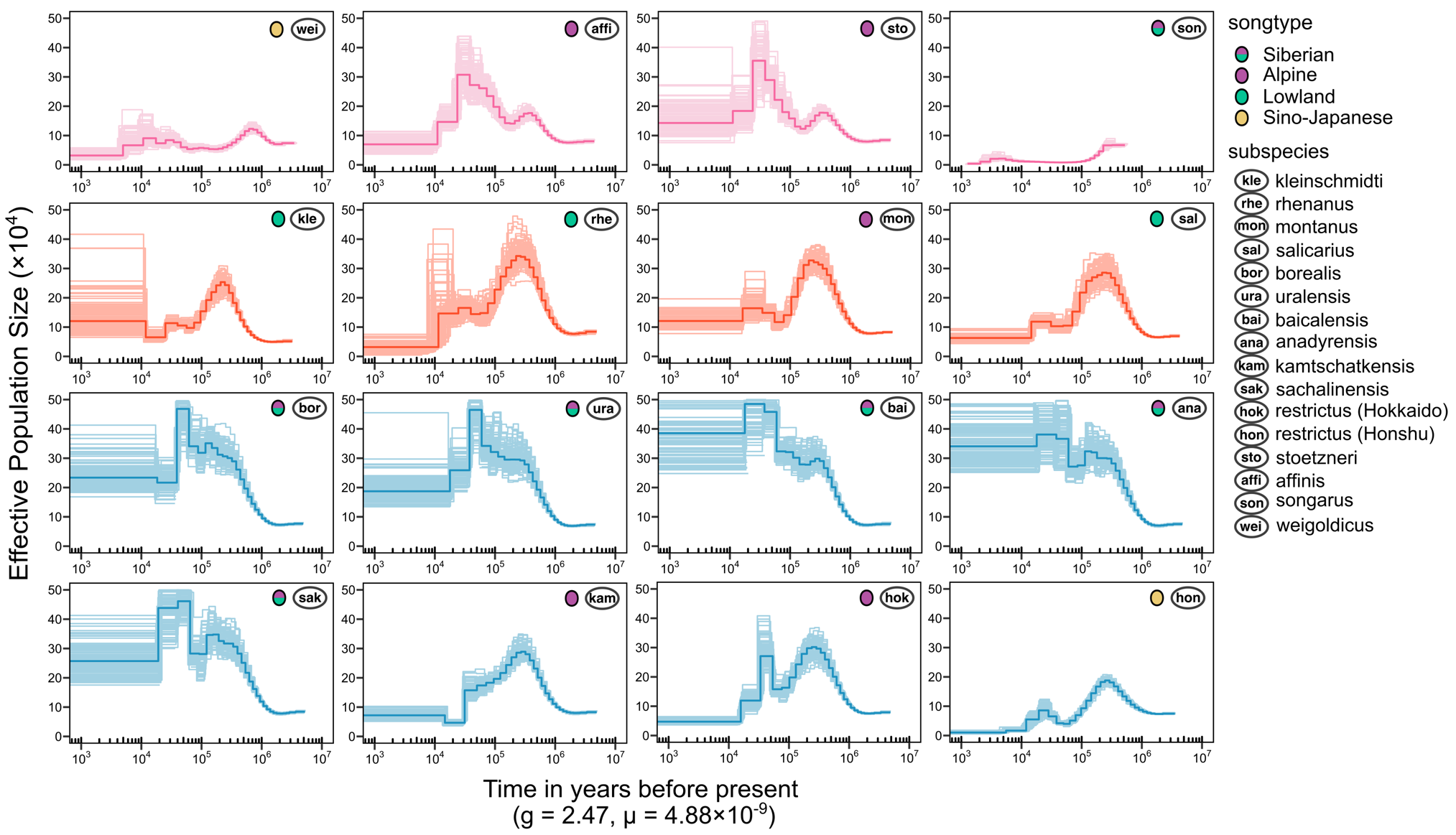

**Supplementary Figure S6.** Population Pairwise Sequentially Markovian Coalescent (PSMC) (Li & Durbin, 2011) demographic reconstruction using autosomal SNPs scaled using a per-site per-generation mutation rate of 4.88 × 10^-9^ from the closely related blue tit (Bergeron et al., 2023) and a generation time of 2.47 years for the willow tit (Bird et al., 2020). The lines with the darker colour represent the medians of 100 replicate runs.

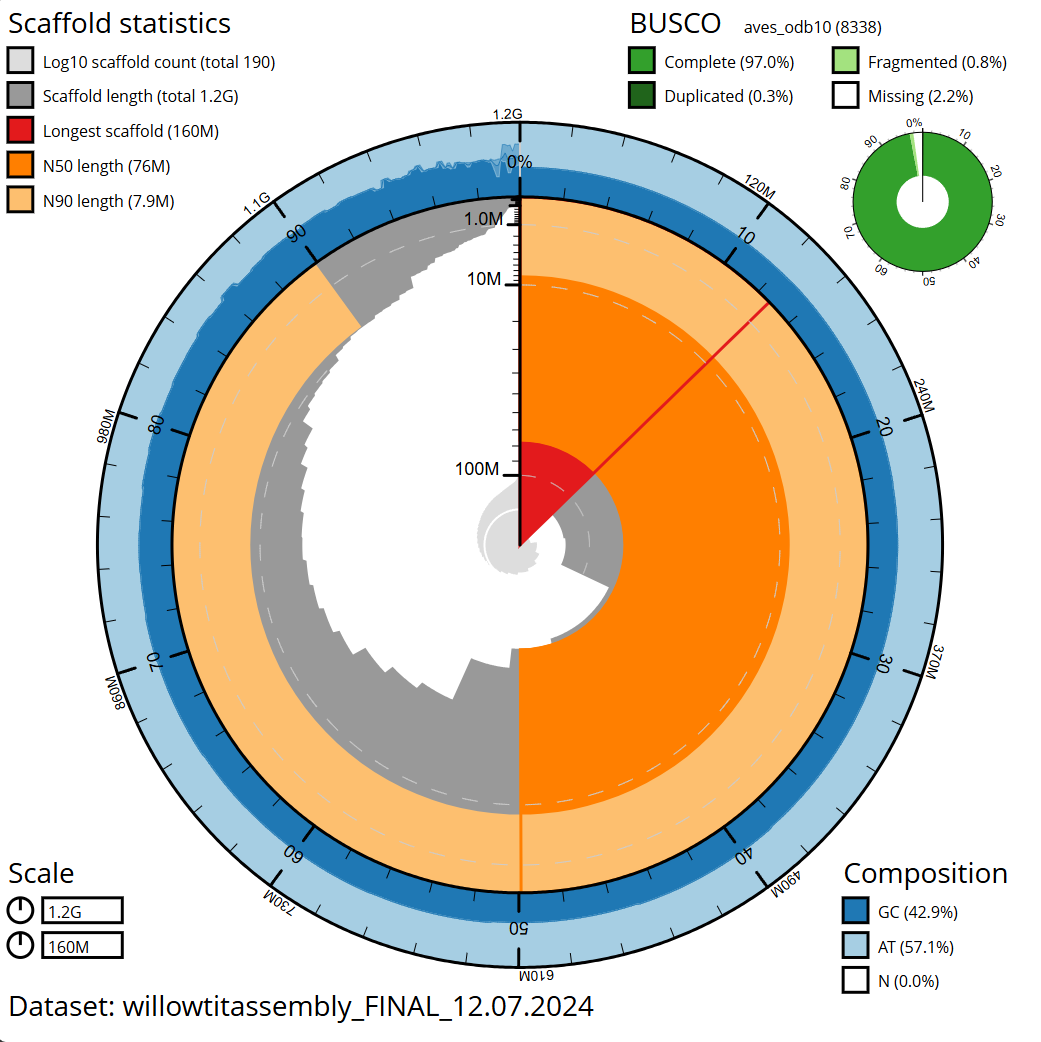

**Supplementary Figure S7.** Snail plot for the willow tit reference genome assembly published in this study ([NCBI GCA_038429875.2](https://www.ncbi.nlm.nih.gov/datasets/genome/GCA_038429875.2/)) made using blobtoolkit v3.2.7 (Laetsch & Blaxter, 2017) showing the reference genome scaffold statistics (top left): scaffold count (191), genome length (1,223,830,793 bp), longest scaffold (156,106,300 bp), N50 length (76,173,615 bp), and N90 length (7,898,092 bp); BUSCO v5.2.2 statistics (top right) (Manni et al., 2021; Simão et al., 2015): complete BUSCOs (8088; 97.0%), complete and single-copy BUSCOs (8067; 96.7%), complete and duplicated BUSCOs (21; 0.3%), fragmented BUSCOs (70; 0.8%), missing BUSCOs (180; 2.2%), and total BUSCOs search (8338) in the aves_odb10 database; and genome composition statistics (bottom right): GC% (42.9%), AT (57.1%). The scale of the snail plot is shown on the bottom left.

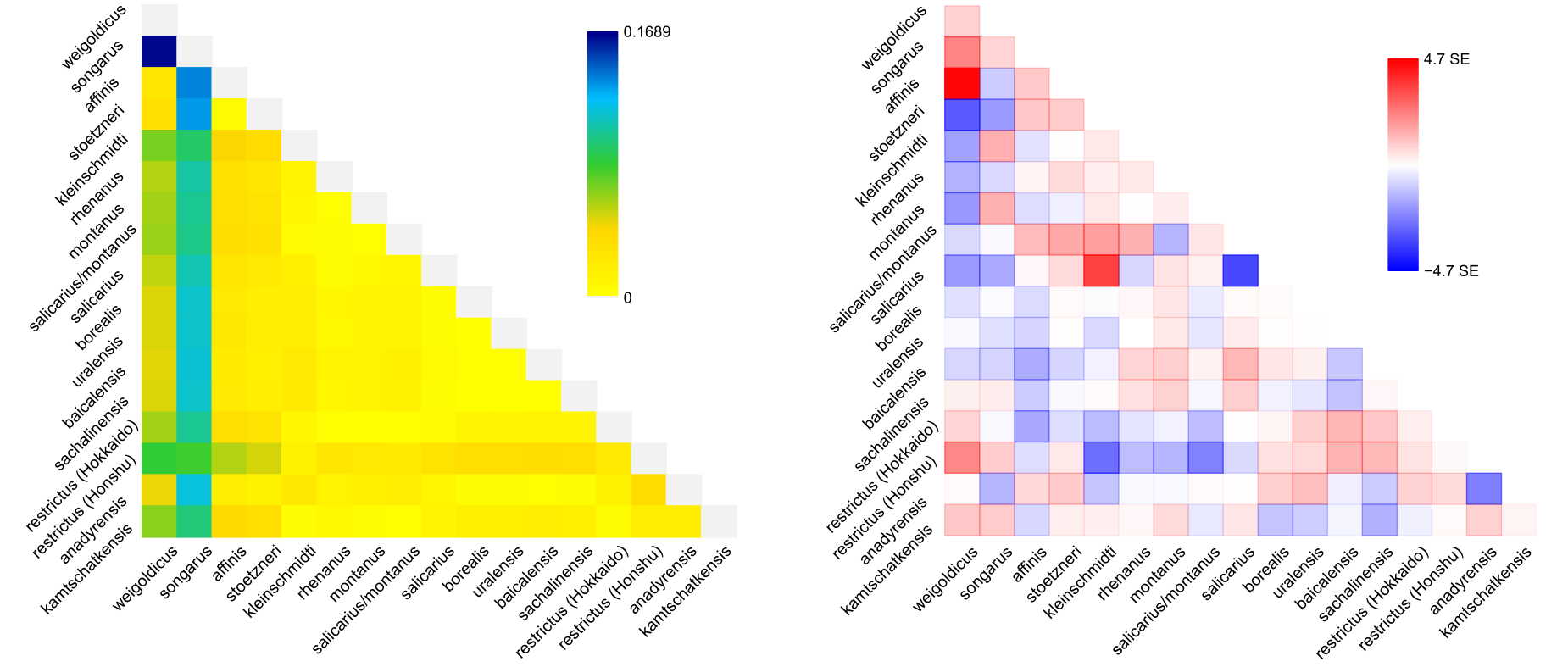

**Supplementary Figure S8.** Drift parameter (left) and the scaled residual fit (right) from the maximum likelihood tree with six migration edges modelled by TreeMix v1.13 (Pickrell & Pritchard, 2012). The model with six migration edges provided an adequate fit to the observed allele-frequency covariance. Deviations exceeding ±3 standard errors are observed between *affinis–weigoldicus, salicarius–kleinschmidti, and salicarius–salicarius,* which represent allele-frequency covariance unexplained by the model. Significant drift was detected within the *songarus* lineage to all willow tit subspecies and *weigoldicus*.

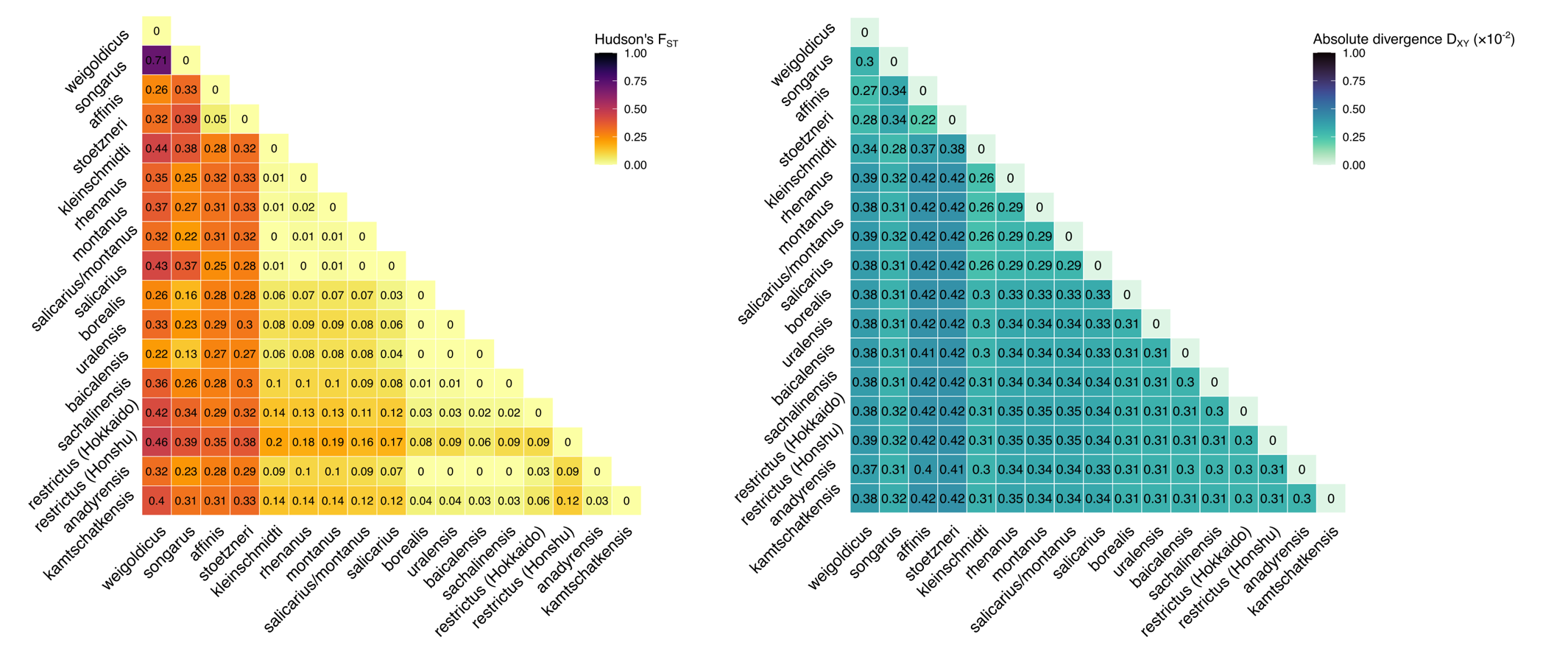

**Supplementary Figure S9.** Genomic differentiation (F_ST_) and absolute divergence (D_XY_) calculated using filtered and masked autosomal SNPs between subspecies: *weigoldicus* (*N* = 3), *affinis* (*N* = 9), *stoetzneri* (*N* = 7), *songarus* (*N* = 2), *kleinschmidti* (*N* = 3), *rhenanus* (*N* = 6), *montanus* (*N* = 5), *salicarius/montanus* (*N* = 7), *salicarius* (*N* = 2), *borealis* (*N* = 10), *uralensis* (*N* = 5), *baicalensis* (*N* = 12), *anadyrensis* (*N* = 2), *kamtschatkensis* (*N* = 4), *sachalinensis* (*N* = 4), *restrictus-*Hokkaido (*N* = 3), *restrictus*-Honshu (*N* = 4). Colours represent genome-wide (autosomal) genomic differentiation (left) and absolute divergence (right, 0–1, lighter colour=less differentiation).

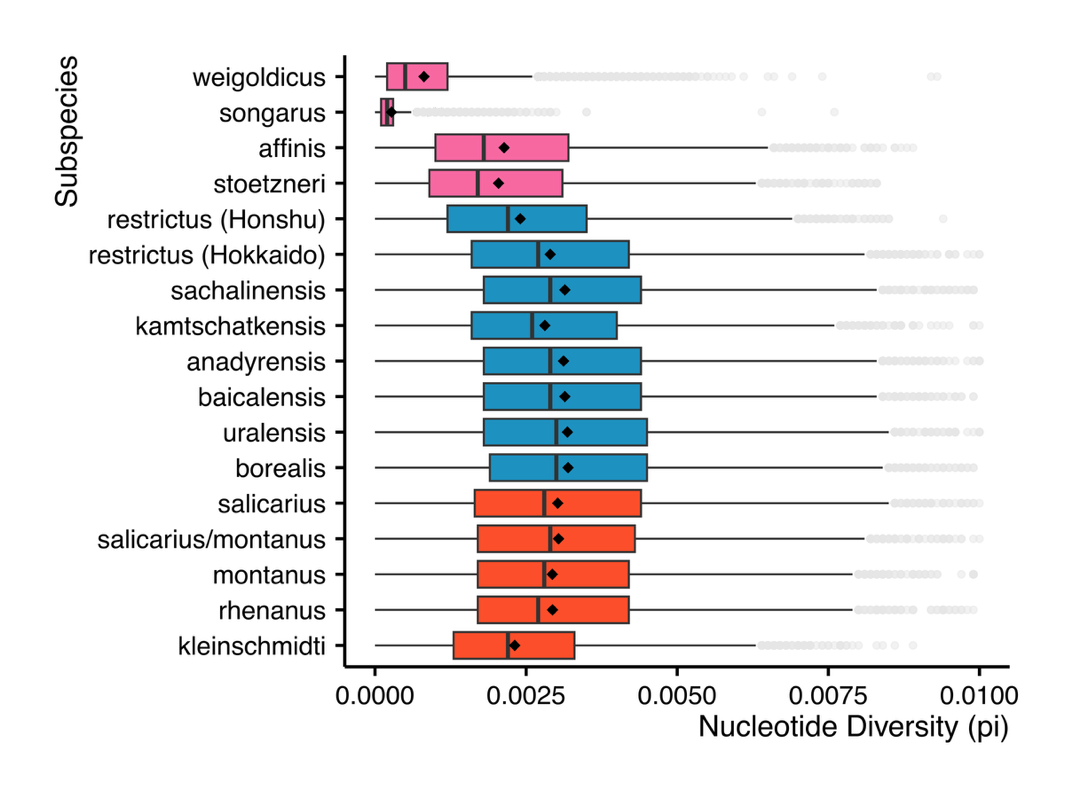

**Supplementary Figure S10.** Box plots of genome-wide nucleotide diversity (π) estimated using filtered and masked autosomal SNPs for each subspecies: *weigoldicus* (*N* = 3), *affinis* (*N* = 9), *stoetzneri* (*N* = 7), *songarus* (*N* = 2), *kleinschmidti* (*N* = 3), *rhenanus* (*N* = 6), *montanus* (*N* = 5), *salicarius/montanus* (*N* = 7), *salicarius* (*N* = 2), *borealis* (*N* = 10), *uralensis* (*N* = 5), *baicalensis* (*N* = 12), *anadyrensis* (*N* = 2), *kamtschatkensis* (*N* = 4), *sachalinensis* (*N* = 4), *restrictus-*Hokkaido (*N* = 3), *restrictus*-Honshu (*N* = 4). The box plots show the interquartile range, the lines represent the median and the diamonds the mean nucleotide diversities of each subspecies.

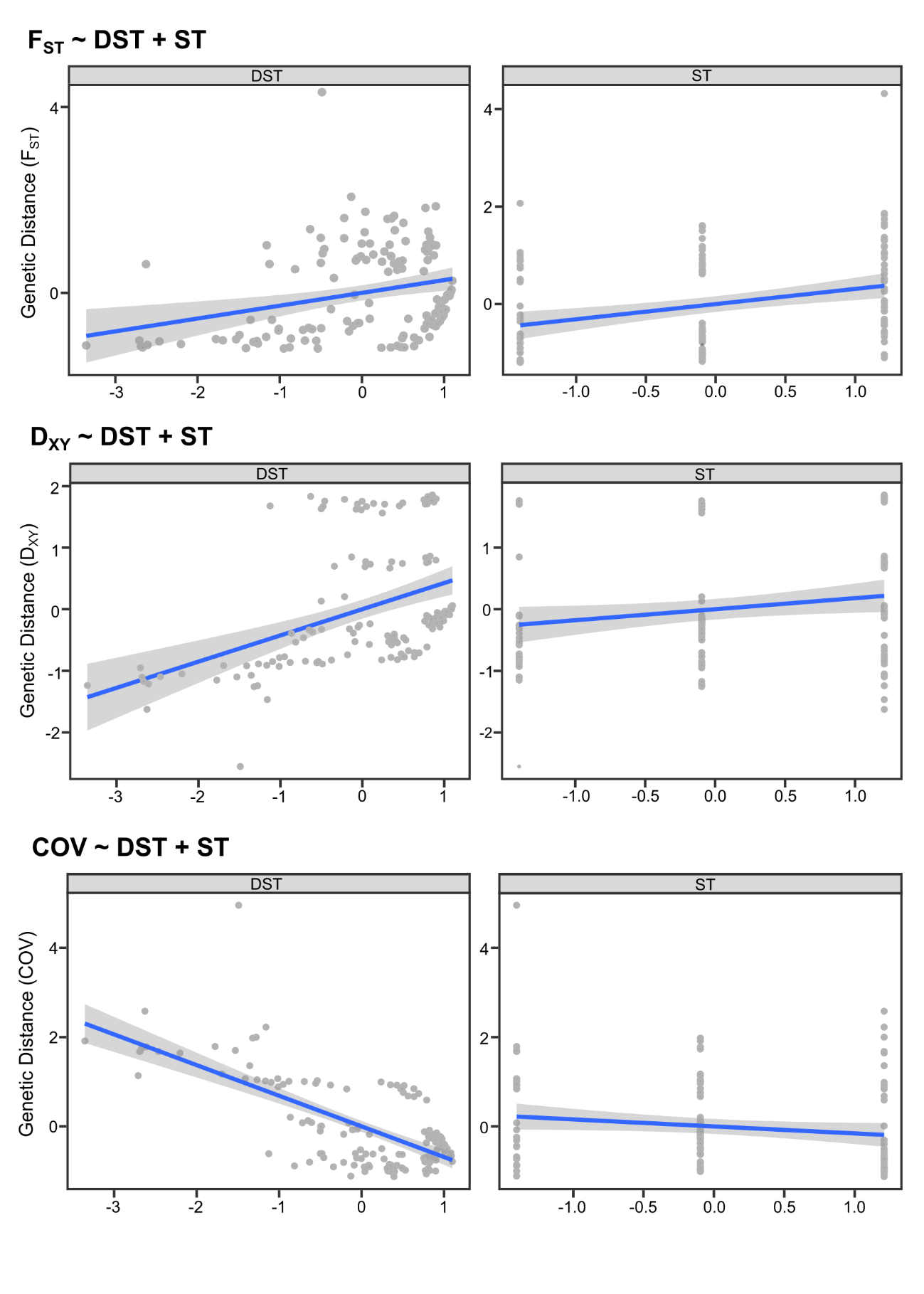

**Supplementary Figure S11.** Multiple regression on distance matrices (MRM) using different genetic distance matrices as the response variable (F_ST_, D_XY_, COV) and geographic distance (DST) and song dissimilarity (ST) as responses.

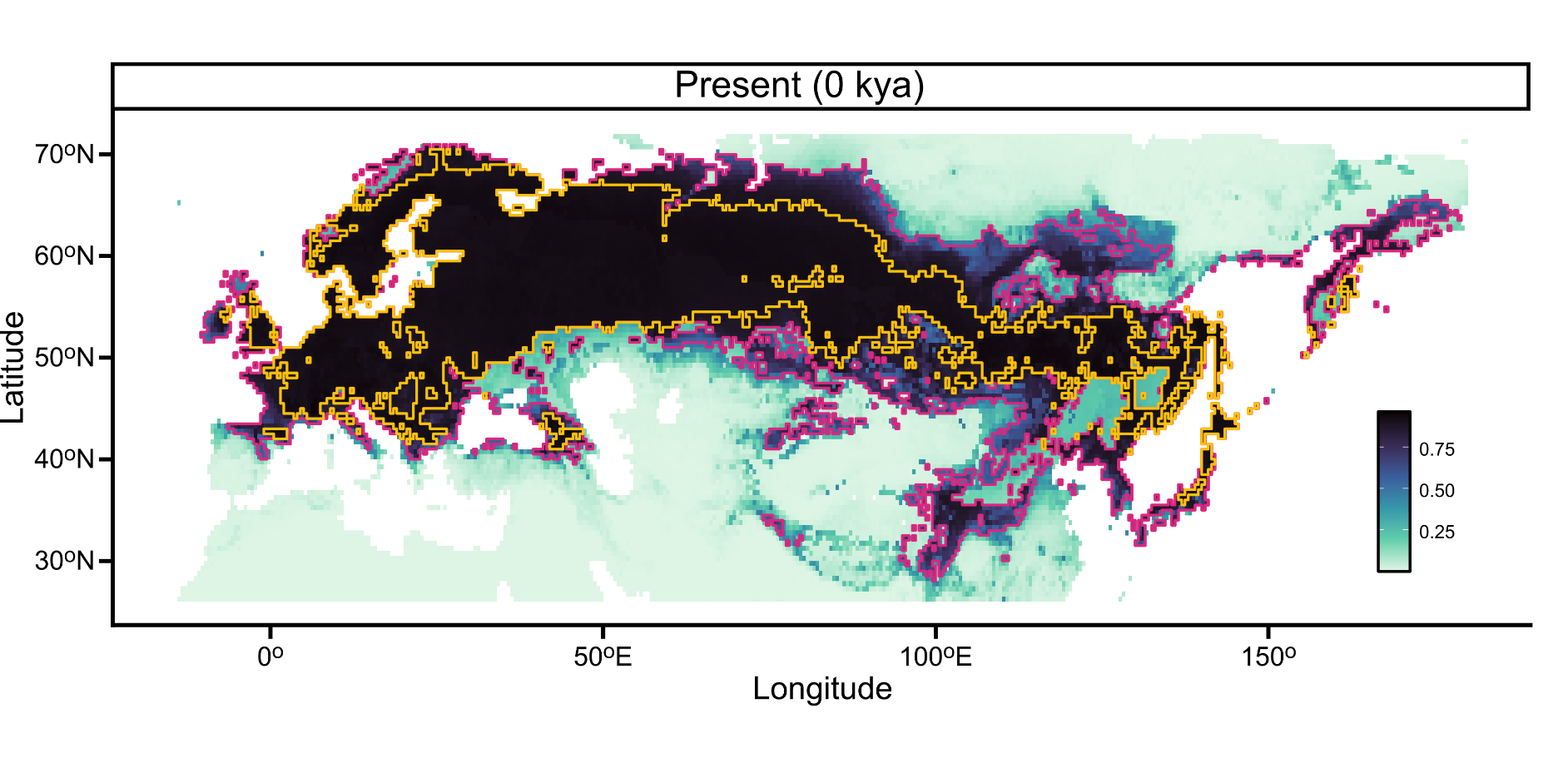

**Supplementary Figure S12.** Ensemble species distribution model (SDM) for the present time (0 kya). The map represents the ensemble model weighted by the area under the receiver operating curve, calculated from 120 species distribution models (30 replicates for each of four modelling algorithms: Maxent, GLM, boosted regression trees, and random forest. We used 9 bioclimatic variables: annual mean temperature (BIO1), temperature seasonality (BIO4), mean temperature of the wettest quarter (BIO8), precipitation of the wettest month (BIO13), precipitation of the driest month (BIO14), precipitation seasonality (BIO15), as well as global biome distribution, annual net primary production, and leaf area index derived from the BIOME4 vegetation model (Kaplan et al., 2003; Krapp et al., 2021). Click [here](https://drive.google.com/file/d/1YJaChcrq1KiXEJxG9dzf5B6JfrtPB8h3/view?usp=sharing) to view the hindcasting from 800–0 kya in 10 ka intervals. Colours represent modelled habitat suitability (0–1; darker colours = higher suitability). The magenta outline represents the area with >0.5 suitability, and the yellow outline shows the area with >0.9 suitability to visualize the fragmentation of suitable habitat.
