## Supplementary Tables for "Cultural transmission and genomic co-divergence in the willow tit across the Palearctic"

**Table S1.** Fitted migration edges from TreeMix with f-branch statistics support

| Donor | Recipient | shared song type | fbC | W | W_JK | SE_JK | p |
|---|---|---|---|---|---|---|---|
| restrictus | stoetzneri | NA | 0.0330801 | 0.111385 | 0.111405 | 0.009706 | $2.225 \times 10^{-308}$ |
| stoetzneri | kleinschmidti | NA | 0.0342743 | 0.042893 | 0.042868 | 0.00389 | $2.225 \times 10^{-308}$ |
| rhenanus | uralensis | Lowland | 0.0439317 | 0.11792 | 0.117903 | 0.003347 | $2.225 \times 10^{-308}$ |
| kleinschmidti | rhenanus | Lowland | 0.0857637 | 0.290935 | 0.291478 | 0.032241 | $2.225 \times 10^{-308}$ |
| montanus | borealis | Alpine | 0.100321 | 0.183985 | 0.1839 | 0.003576 | $2.225 \times 10^{-308}$ |
| kamtschatkensis | anadyrensis | Alpine | 0.115185 | 0.362824 | 0.36273 | 0.034 | $2.225 \times 10^{-308}$ |

fbC = f-branch estimates, W = weight of migration edge, W_JK = jack-knife estimate of weight, SE_JK = standard error of the jack-knife estimate, p = p-value of weight estimate

**Table S2.** Multiple Regression on Multiple Distance Matrices model of Isolation-by-Distance (IBD) and Isolation-by-Song (IBS)

| Model | $R^2$ | F | **p (model)** | Predictor | ß coefficient | **p** (N perm=10000) | 95% Lower | 95% Upper |
|---|---|---|---|---|---|---|---|---|
| **FST ~ DST + ST** | 0.166212 | 13.25648 | 0.009799 | DST | 2.61E-01 | 0.00009999 | 0.1291129 | 0.3932427 |
| | | | | **ST** | **2.84E-01** | **0.049995** | 0.1517778 | 0.4159076 |
| **DXY ~ DST + ST** | 0.177846 | 14.38506 | 8.50E-03 | DST | 3.97E-01 | 0.00009999 | 0.26584181 | 0.5281224 |
| | | | | **ST** | **1.03E-01** | **0.35056494** | -0.0280489 | 0.2342317 |
| **COV ~ DST + ST** | 0.47729 | 60.72158 | 1.00E-04 | DST | -6.77E-01 | 0.00009999 | -0.7820503 | -0.5729186 |
| | | | | **ST** | **-7.73E-02** | **0.21307869** | -0.1819152 | 0.02721655 |

FST=F-statistics, DXY=absolute divergence, COV=covariance matrix as implemented in TreeMix

**Table S3.** Sample location, molecular-typed sex, inferred subspecies, song type, locality, provider, and coverage.

| SampleID | Inferred Subspecies | Song type | Sex | Lat | Long | Locality | Provider | Coverage (G=1.2Gb) |
|---|---|---|---|---|---|---|---|---|
| M1864_2 | P. weigoldicus | Sino-Japanese | M | 33.22 | 103.73 | Min Shan, kleine Forststation, China | JM | 16.72 |
| M1865_2 | P. weigoldicus | Sino-Japanese | M | 33.22 | 103.73 | Min Shan, kleine Forststation, China | JM | 17.35 |
| M5672 | P. m. rhenanus | Lowland | M | 49.75 | 8.37 | Constant Effort Site near Eich am Rhein, Germany | JM | 17.83 |
| MTDC48361 | P. m. songarus | Siberian | M | 42.65 | 76.99 | Issyk-Kul, Kyrgyzstan | JM | 16.63 |
| N_102 | P. m. salicarius / montanus | Alpine/Lowland | M | 48.78 | 13.95 | Nová Pec, SSE Volary, Czech Republic | JM | 16.28 |
| N_1085 | P. m. anadyrensis | Siberian | M | 64.67 | 170.42 | Markowo, Russia | JM | 17.04 |
| N_1156 | P. m. baicalensis | Siberian | F | 46.52 | 136.95 | Primorskij Krai, Ochotnicij, Russia | JM | 16.83 |
| N_1157 | P. m. baicalensis | Siberian | M | 46.52 | 136.95 | Primorskij Krai, Ochotnicij, Russia | JM | 17.27 |
| N_1158 | P. m. baicalensis | Siberian | F | 46.52 | 136.95 | Primorskij Krai, Ochotnicij, Russia | JM | 16.84 |
| N_1344 | P. m. baicalensis | Siberian | M | 49.00 | 131.72 | Birakan, Russia | JM | 17.13 |
| N_1347 | P. m. baicalensis | Siberian | F | 49.00 | 131.72 | Birakan, Russia | JM | 16.73 |
| N_1354 | P. m. baicalensis | Siberian | M | 49.00 | 131.72 | Birakan, Russia | JM | 16.13 |
| N_1355 | P. m. baicalensis | Siberian | F | 49.00 | 131.72 | Birakan, Russia | JM | 16.71 |
| N_1384 | P. m. baicalensis | Siberian | M | 43.67 | 134.17 | Ussuri River, Russia | JM | 16.98 |
| N_1386 | P. m. baicalensis | Siberian | M | 43.67 | 134.17 | Ussuri River, Russia | JM | 16.91 |
| N_1390 | P. m. baicalensis | Siberian | M | 43.67 | 134.17 | Ussuri River, Russia | JM | 16.83 |
| N_1409 | P. m. sachalinensis | Siberian | M | 46.97 | 142.22 | Chaplanovo; 16 km SW Kholmsk, Russia | JM | 18.04 |
| N_1411 | P. m. sachalinensis | Siberian | M | 46.97 | 142.22 | Chaplanovo; 16 km SW Kholmsk, Russia | JM | 21.48 |
| N_1413 | P. m. sachalinensis | Siberian | M | 46.97 | 142.22 | Chaplanovo; 16 km SW Kholmsk, Russia | JM | 16.06 |

| | | | | | | | | |
|---|---|---|---|---|---|---|---|---|
| N_1414 | P. m. sachalinensis | Siberian | M | 46.97 | 142.22 | Chaplanovo; 16 km SW Kholmsk, Russia | JM | 17.15 |
| N_1609 | P. m. songarus | Siberian | M | 43.77 | 77.12 | Almaatinka valley, Kazakhstan | JM | 16.98 |
| N_1870 | P. m. affinis | Alpine | F | 34.92 | 103.72 | Lianhua Shan Nature Reserve, Shahetan, China | JM | 16.48 |
| N_1913 | P. m. salicarius / montanus | Alpine/Lowland | M | 48.78 | 13.93 | Nová Pec, obere Moldau, Czech Republic | JM | 16.96 |
| N_1927 | P. m. salicarius / montanus | Alpine/Lowland | M | 48.78 | 13.93 | Nová Pec, obere Moldau, Czech Republic | JM | 17.05 |
| N_1934_1_1_368 | P. m. kleinschmidti | Lowland | M | 55.79 | -4.09 | Blantyre, South Lanarkshire, Scotland | NHM | 18.92 |
| N_1934_1_1_5973 | P. m. kleinschmidti | Lowland | M | 51.81 | -0.69 | Wilstone Reservoirs, Tring, Hertfordshire, England | NHM | 17.17 |
| N_1951_13_1951 | P. m. kleinschmidti | Lowland | M | 51.40 | -1.20 | Midgham, Berkshire, England | NHM | 16.67 |
| YOI-N_1992_0123 | P. m. restrictus (Honshu) | Sino-Japanese | M | 35.31 | 138.93 | Gotemba, Shizuoka, Japan | YOI | 16.72 |
| YOI-N_1998_0130 | P. m. restrictus (Honshu) | Sino-Japanese | F | 32.98 | 130.81 | Kikuchi, Kumamoto, Japan | YOI | 16.84 |
| YOI-N_2001_0025 | P. m. restrictus (Hokkaido) | Alpine | F | 43.05 | 141.32 | Chuo-ku, Sapporo, Hokkaido, Japan | YOI | 16.93 |
| YOI-N_2001_0205 | P. m. restrictus (Honshu) | Sino-Japanese | F | 35.31 | 138.93 | Gotemba, Shizuoka, Japan | YOI | 16.84 |
| YOI-N_2002_0292 | P. m. restrictus (Hokkaido) | Alpine | F | 43.33 | 145.58 | Nemuro, Hokkaido, Japan | YOI | 17.92 |
| YOI-N_2003_5409 | P. m. restrictus (Honshu) | Sino-Japanese | M | 35.41 | 138.86 | Yamanakako-mura, Minami-tsuru-gun, Yamanashi, Japan | YOI | 16.72 |
| N_2004 | P. m. salicarius / montanus | Alpine/Lowland | M | 48.10 | 11.35 | Pionier Area, Krailling, Germany | UK | 17.01 |
| YOI-N_2010_1759 | P. m. restrictus (Hokkaido) | Alpine | F | 45.12 | 142.36 | Hamatombetsu-cho, Esashi-gun, Hokkaido, Japan | YOI | 20.06 |
| N_21 | P. m. rhenanus | Lowland | F | 49.97 | 8.12 | Wackernheim, bei Mainz, Germany | JM | 16.29 |
| N_351 | P. m. affinis | Alpine | M | 36.93 | 102.65 | Bei Shan, Chingshiling Reserve, China | JM | 16.99 |
| N_355 | P. m. affinis | Alpine | M | 36.93 | 102.65 | Bei Shan, Chingshiling Reserve, China | JM | 16.93 |
| N_401 | P. m. affinis | Alpine | M | 36.93 | 102.65 | Bei Shan, Chingshiling Reserve, China | JM | 17.08 |
| N_403 | P. m. affinis | Alpine | F | 36.93 | 102.65 | Bei Shan, Chingshiling Reserve, China | JM | 16.85 |

| | | | | | | | | |
|---|---|---|---|---|---|---|---|---|
| N_4843 | P. m. kamtschatkensis | Alpine | F | 53.18 | 158.38 | Elisodo town, SE Kamtchatka, Russia | JM | 17.03 |
| N_4844 | P. m. kamtschatkensis | Alpine | F | 53.18 | 158.38 | Elisodo town, SE Kamtchatka, Russia | JM | 17.11 |
| N_4846 | P. m. kamtschatkensis | Alpine | F | 53.18 | 158.38 | Elisodo town, SE Kamtchatka, Russia | JM | 16.67 |
| N_4847 | P. m. kamtschatkensis | Alpine | M | 53.18 | 158.38 | Elisodo town, SE Kamtchatka, Russia | JM | 19.01 |
| N_5056 | P. m. montanus | Alpine | F | 46.52 | 11.50 | Völser Aicha bei Völs am Schlern, Austria | JM | 18.34 |
| N_5057 | P. m. montanus | Alpine | M | 46.52 | 11.50 | Völser Aicha bei Völs am Schlern, Austria | JM | 17.64 |
| N_5129 | P. m. rhenanus | Lowland | M | 49.75 | 8.37 | Constant Effort Site near Eich am Rhein, Germany | JM | 17.67 |
| N_6 | P. m. rhenanus | Lowland | F | 49.95 | 8.17 | Oberolmer Wald bei Mainz, Germany | JM | 16.78 |
| N_6489 | P. m. stoetzneri | Alpine | M | 38.78 | 105.85 | Helan Shan, Gun Zhong Kou, holiday resort, China | JM | 16.96 |
| N_6490 | P. m. stoetzneri | Alpine | M | 38.78 | 105.85 | Helan Shan, Gun Zhong Kou, holiday resort, China | JM | 16.83 |
| N_685 | P. m. baicalensis | Alpine/Lowland /Sino-Japanese | F | 50.12 | 88.35 | Tschagan-Uzun, Russia | JM | 16.77 |
| N_689 | P. m. montanus | Alpine | M | 47.37 | 13.12 | Mühlbach am Hochkönig, Kopphütte, Austria | JM | 16.28 |
| N_690 | P. m. montanus | Alpine | F | 47.37 | 13.12 | Mühlbach am Hochkönig, Kopphütte, Austria | JM | 16.76 |
| N_6926 | P. m. affinis | Alpine | M | 34.92 | 103.72 | Lianhua Shan Nature Reserve, Shahetan, China | JM | 16.18 |
| N_6928 | P. m. affinis | Alpine | M | 34.92 | 103.72 | Lianhua Shan Nature Reserve, Shahetan, China | JM | 16.7 |
| N_6932 | P. m. affinis | Alpine | F | 34.92 | 103.72 | Lianhua Shan Nature Reserve, Shahetan, China | JM | 16.93 |
| N_7 | P. m. rhenanus | Lowland | M | 49.95 | 8.17 | Oberolmer Wald bei Mainz, Germany | JM | 16.66 |
| N_7083 | P. m. rhenanus | Lowland | F | 49.75 | 8.37 | Constant Effort Site near Eich am Rhein, Germany | JM | 16.92 |
| N_79 | P. m. montanus | Alpine | F | 46.72 | 11.65 | Bressanone, Italy | JM | 16.37 |
| N_8105 | P. m. affinis | Alpine | M | 34.92 | 103.72 | Lianhua Shan Nature Reserve, Shahetan, China | JM | 16.81 |
| N_8171 | P. weigoldicus | Sino-Japanese | F | 31.87 | 96.55 | San Jiang Yuan National Reserve, China | JM | 16.7 |

| | | | | | | | | |
|---|---|---|---|---|---|---|---|---|
| N_83 | P. m. salicarius / montanus | Alpine/Lowland | F | 48.78 | 13.95 | Nová Pec, SSE Volary, Czech Republic | JM | 17.56 |
| N_8564 | P. m. stoetzneri | Alpine | F | 39.82 | 115.57 | Baicaopan Geopark, China | JM | 19.03 |
| N_8565 | P. m. stoetzneri | Alpine | M | 39.82 | 115.57 | Baicaopan Geopark, China | JM | 19.12 |
| N_8583 | P. m. stoetzneri | Alpine | F | 40.58 | 117.48 | Wuling Shan NE Beijing, China | JM | 18.47 |
| N_8585 | P. m. stoetzneri | Alpine | F | 40.58 | 117.48 | Wuling Shan NE Beijing, China | JM | 18.1 |
| N_8586 | P. m. stoetzneri | Alpine | F | 40.58 | 117.48 | Wuling Shan NE Beijing, China | JM | 17.42 |
| N_9104 | P. m. salicarius | Lowland | F | 51.30 | 14.90 | Horka Johannenhof, Germany | JM | 16.13 |
| N_94 | P. m. salicarius / montanus | Alpine/Lowland | M | 48.78 | 13.95 | Nová Pec, SSE Volary, Czech Republic | JM | 17.13 |
| N_9449 | P. m. baicalensis | Siberian | F | 50.17 | 106.17 | Schaamar, Mongolia | JM | 16.78 |
| N2005 | P. m. salicarius / montanus | Alpine/Lowland | F | 48.10 | 11.35 | Pionier Area, Krailling, Germany | UK | 53.06 |
| T1481 | P. m. borealis | Siberian | M | 65.01 | 25.47 | Oulu, Finland | LK | 16.51 |
| T1483 | P. m. borealis | Siberian | M | 65.01 | 25.47 | Oulu, Finland | LK | 18.77 |
| T178 | P. m. uralensis | Siberian | M | 56.43 | 61.92 | Kamensk Uralski, Russia | LK | 17.13 |
| T179 | P. m. uralensis | Siberian | F | 56.43 | 61.92 | Kamensk Uralski, Russia | LK | 16.27 |
| T180 | P. m. uralensis | Siberian | M | 56.43 | 61.92 | Kamensk Uralski, Russia | LK | 16.98 |
| T181 | P. m. uralensis | Siberian | M | 56.43 | 61.92 | Kamensk Uralski, Russia | LK | 16.67 |
| T185 | P. m. uralensis | Siberian | F | 56.43 | 61.92 | Kamensk Uralski, Russia | LK | 16.95 |
| T186 | P. m. borealis | Siberian | M | 60.74 | 46.37 | Kuzino, Russia | LK | 16.76 |
| T187 | P. m. borealis | Siberian | F | 60.74 | 46.37 | Kuzino, Russia | LK | 16.77 |
| T188 | P. m. borealis | Siberian | M | 60.74 | 46.37 | Kuzino, Russia | LK | 17.18 |
| T189 | P. m. borealis | Siberian | M | 60.74 | 46.37 | Kuzino, Russia | LK | 18.9 |
| T190 | P. m. borealis | Siberian | M | 60.74 | 46.37 | Kuzino, Russia | LK | 17.24 |
| T192 | P. m. anadyrensis | Siberian | F | 59.56 | 150.81 | Magadan, Russia | LK | 16.6 |
| T193 | P. m. anadyrensis | Siberian | F | 59.56 | 150.81 | Magadan, Russia | LK | 17.39 |
| T194 | P. m. anadyrensis | Siberian | F | 59.56 | 150.81 | Magadan, Russia | LK | 16.99 |
| T196 | P. m. anadyrensis | Siberian | M | 59.56 | 150.81 | Magadan, Russia | LK | 17.16 |

| | | | | | | | | |
|---|---|---|---|---|---|---|---|---|
| T248 | P. m. baicalensis | Siberian | F | 41.80 | 128.32 | Samjiyon, North Korea | LK | 16.93 |
| T37 | P. m. borealis | Siberian | M | 58.95 | 17.15 | Tovetorp, Sweden | LK | 16.67 |
| T72 | P. m. borealis | Siberian | F | 58.95 | 17.15 | Tovetorp, Sweden | LK | 16.88 |
| T79 | P. m. salicarius | Lowland | F | 52.38 | 9.73 | Hannover, Germany | LK | 17.17 |

JM= Jochen Martens / Senckenberg Natural History Collections Dresden; NHM= Natural History Museum; YOI= Yamashina Institute for Ornithology; UK=Ulrich Knief, University of Freiburg; Laura Kvist, University of Oulu

**Table S4.** Final reference genome assembly metrics

| **Assembly metrics** | |
|---|---|
| n contigs >= 10000 bp | 188 |
| n contigs >= 25000 bp | 178 |
| n contigs >= 50000 bp | 151 |
| Total length >= 10000 bp | 1223820793 |
| Total length >= 25000 bp | 1223616065 |
| Total length >= 50000 bp | 1222626571 |
| n contigs | 191 |
| Largest contig (bp) | 156106300 |
| Total length (bp) | 1223830793 |
| GC (%) | 42.94 |
| N50 | 76173615 |
| N75 | 21566885 |
| L50 | 6 |
| L75 | 15 |
| **BUSCO (aves_odb10, n=8338)** | |
| Complete BUSCOs | 97% (n=8088) |
| Complete and single-copy BUSCOs | 96.7% (n=8067) |
| Complete and duplicated BUSCOs | 0.3% (n=21) |
| Fragmented BUSCOs | 0.8% (n=70) |
| Missing BUSCOs | 2.2% (n=180) |

**Table S5.** Species Distribution Models and hindcasting evaluation. The reported Area under the curve (AUC) is the average for the 30 model replicates run for each algorithm.

| Models | AUC (mean ± SD) |
|---|---|
| Full ensemble model (n = 30) | 0.997 ± 0.0016 |
| Random Forest (n = 30) | 0.995 ± 0.0047 |
| Generalized Linear Model (n = 30) | 0.979 ± 0.0344 |
| Boosted Regression Trees (n = 30) | 0.997 ± 0.0016 |
| Maximum Entropy (n = 30) | 0.994 ± 0.0014 |
